# High-latitude marginal reefs support fewer but bigger corals than their tropical counterparts

**DOI:** 10.1101/2022.10.20.513025

**Authors:** Fiona Chong, Brigitte Sommer, Georgia Stant, Nina Verano, James Cant, Liam Lachs, Magnus L Johnson, Daniel R Parsons, John M Pandolfi, Roberto Salguero-Gómez, Matthew Spencer, Maria Beger

## Abstract

**Aim:** Despite the awareness that climate change impacts are typically detrimental to tropical coral reefs, the effect of increasing environmental stress and variability on the population size structure of coral species remains poorly understood. This gap in knowledge limits our ability to effectively conserve coral reef ecosystems because size specific dynamics are rarely incorporated. Our aim is to quantify variation in the size structure of coral populations along a tropical-to-subtropical environmental gradient.

**Location:** 20 coral populations along a latitudinal gradient on the east coast of Australia (∼23°S to 30°S).

**Time Period:** Between 2010 and 2018.

**Major taxa studied:** Scleractinian corals.

**Methods:** We apply two methods to quantify the relationship between environmental covariates and coral population size structure along a latitudinal environmental gradient. First, we use linear regression with summary statistics, such as median size as response variables; a method frequently favoured by ecologists. The second method is compositional functional regression, a novel method using entire size-frequency distributions as response variables. We then predict coral population size structure with increasing environmental stress and variability.

**Results:** Compared to tropical reefs, we find fewer but larger coral colonies in marginal reefs, where environmental conditions are more variable and stressful for corals in the former. Our model predicts that coral populations may become gradually dominated by larger colonies (> 148 cm^2^) with increasing environmental stress.

**Main conclusions:** With increasing environmental stress and variability, we can expect shifts in coral population size structure towards more larger colonies. Fewer but bigger corals suggest low survival, slow growth, and poor recruitment. This finding is concerning for the future of coral reefs as it implies populations may have low recovery potential from disturbances. We highlight the importance and usefulness of continuously monitoring changes to population structure over large spatial scales.

**Data availability:** Data is supplied in the supplementary information, or upon request. Once accepted for publication it will be made openly available on Dryad.

## Introduction

Population size has been a primary metric of population persistence and viability for decades (Shaffer, 1981; Dietzel *et al*., 2021). However, the size structure of a population (*i*.*e*., how many individuals of a given size range there are in the population) is as important, if not more so, for determining persistence and viability, especially in slow growing, sessile organisms (*e*.*g*., McClanahan *et al*., 2008; Riegl *et al*., 2012; Cousins *et al*., 2014). The structure of a population details important features regarding individual heterogeneity that ultimately predict population outcomes better than simply population size (Hunter *et al*., 2010; Radchuk *et al*., 2013). Consequently, in recent decades, population structure has become the focus of demographic models (Easterling *et al*., 2000; Caswell, 2001; Merow *et al*., 2014).

External (a)biotic factors such as climate change (*e*.*g*., Radchuk *et al*., 2013; Vetter *et al*., 2020) can lead to shifts in population structure when the underlying vital rates (*e*.*g*., survival, change in size, reproduction) are affected differently. For example, Radchuk *et al*. (2013) showed that increases in temperature improve the fecundity of female bog fritillary butterflies (*Boloria eunomia*) and the survival of most life stages, except for the overwintering larvae. Yet the viability of the butterfly population is highly sensitive to the survival of overwintering larvae (Radchuk *et al*., 2013), meaning that low larval survival, as a result of warming, would be detrimental to the viability of this population. However, warming is not constant, and is only one of many aspects of climate change (Dixon *et al*., 2021), to which species and population responses are complex and not well understood (Lawson *et al*., 2015; Tavecchia *et al*., 2016). Further, creating meaningful and realistic experimental manipulations to understand future climate effects on population structure might be impracticable (Kreyling *et al*., 2014). An alternative approach to understand the effect of climate change on populations is to sample from natural populations exposed to different environmental conditions (shift in mean conditions, increased variability and extremes) (Kreyling *et al*., 2014), *e*.*g*., at the biogeographic scale. Despite the potential of this approach for predicting the effects of climate change on population viability, we know of no such studies.

Coral reefs are challenged by many anthropogenic perturbations, with climate change remains the dominant threat (Pandolfi, 2015; Hoegh-Guldberg *et al*., 2017; Hughes *et al*., 2017). Climate change will likely increase thermal stress (Dixon *et al*., 2022), flooding (Vitousek *et al*., 2017) and storm intensity (Reguero *et al*., 2019).

These disturbances directly and indirectly induce coral mortality and changes in community composition (Hughes *et al*., 2012; Ceccarelli *et al*., 2020; Brunner *et al*., 2021) and coral population size structure (*e*.*g*., Hughes *et al*., 2018; Pisapia *et al*., 2019; Dietzel *et al*., 2020; Lachs *et al*., 2021). Considering that the vital rates of survival, growth, and reproduction follow consistent allometric scaling in corals (Dornelas *et al*., 2017; Madin *et al*., 2020), changes to coral population size structure will have major consequences for their dynamics and viability. Indeed, small corals tend to have a higher probability of whole-colony mortality, while larger corals have higher partial mortality (*i*.*e*., shrinkage) and fission (Hughes & Connell, 1987; Hughes & Tanner, 2000; Madin *et al*., 2020). Large corals also have higher reproduction, but lower relative growth rates (Connell, 1973; Dornelas *et al*., 2017). Because of these allometric relationships, investigating differences in size structure across populations experiencing increased disturbance can reveal the underlying ecological mechanisms that drive population viability.

Previous studies have examined changes in coral population size structure using summary statistics such as mean size, variance, skewness, and kurtosis (*e*.*g*., Bak & Meesters, 1998; Anderson & Pratchett, 2014). These metrics characterize aspects of the shape of the size-frequency distribution. However, the summary statistics approach involves making arbitrary choices about which statistics to include, and does not use all the information in the distribution (Talská *et al*., 2018). Also, the ecological interpretation of measures such as kurtosis is not straightforward.

Adjeroud *et al*. (2007) observed negative kurtosis (a flattened distribution, with a wide peak around the mean) for a fast-growing species, and the opposite for a slow-growing species. Since then, coral reef ecologists have related this metric to population growth and turnover rates, but the conditions under which the proposed relationship between kurtosis and growth rate holds are unclear. The assessment and comparison of entire coral size-frequency distributions as probability density functions can overcome these challenges. Recent advances in functional data analysis (Ramsay *et al*., 2009; Talská *et al*., 2018) remove the need to arbitrarily select a few summary statistics as response variables. Since the entire probability density function is treated as the response variable (Talská *et al*., 2018), the method can accurately quantify which coral sizes are most affected by the explanatory variables. This approach is likely to better capture the effects of environmental stress on coral size-frequency distributions, allowing for improved comparisons and understanding of their dynamics under stress.

Here, we examine the effect of environmental stress on scleractinian coral population size structure in eastern Australia. Using a space-for-time approach (Kreyling *et al*., 2014) that substitutes a tropical-to-subtropical environmental gradient with future environmental change, we aim to understand how reef population dynamics responds to climate change. We use two methodologies: 1) linear regression with summary statistics as response variables, an approach classically favoured by coral reef ecologists, and 2) a novel compositional functional regression approach (Talská *et al*., 2018) that has not been used in this context despite its potential. At higher latitudes, where conditions are harsh due to extremes in temperature, light levels and storm events, we expect fewer small coral colonies, because coral mortality rates are generally highest for the smallest corals (Connell, 1973), and sexual recruitment rates are low (Harriott & Banks, 1995; Abrego *et al*., 2021; Cant *et al*., 2022). Potential differences in population size structure of corals along this environmental gradient might indicate the effect of stress on coral population dynamics, providing a lens to the future, where reefs might be affected by increased disturbances as a result of climate change.

## Methods

### Data collection

The eastern Australian biogeographic transition zone is a unique region in which to observe coral population dynamics. There, coral communities occur from tropical Queensland’s Great Barrier Reef (GBR) to the temperate, sometimes kelp-dominated rocky reefs in New South Wales. With increasing latitude, sea surface temperature and incident light intensity decline, while storm intensity and frequency increase (Pepler & Coutts-Smith, 2013), making the reef habitat increasingly marginal for tropical hard corals (Harriott & Smith, 2000; Sommer *et al*., 2018). Multiple oceanographic currents are present in the region, with the Eastern Australian Current (EAC) being the largest (Baird *et al*., 2008). The EAC runs approximately 50 km offshore (Malcolm *et al*., 2011), transporting warm, tropical waters from the Coral Sea poleward. The current may also be a source of fresh genetic material for the downstream reefs (Beger *et al*., 2014; Sommer *et al*., 2014). Although a recent study has suggested that coral larvae dispersed from the southern GBR have a low probability of being received at higher latitude reefs (Mizerek *et al*., 2021), where endemic coral species are increasingly found (*e*.*g*., Schmidt-Roach et al., 2013; Baird et al., 2017). The eastern Australian biogeographic transition zone thus represents a natural laboratory that allows the examination of differences in coral population size structure in response to increasing environmental stress.

We sampled 20 coral populations in the eastern Australian biogeographic transition zone using underwater photographic benthic transect surveys. 12 sites were sampled in September 2018, while the eight other sites were sampled in either 2010, 2011, 2012 or 2016 (Figure 1; Table S1). At each site, three 30 m belt transects were haphazardly run at 8-10 m water depth. Downward-facing photographs were taken every metre, from approximately 70 cm above the benthos. Each included a 50 cm calibration stick held at the level of the substrate (as in Sommer *et al*., 2011). Two cameras were used: a Canon S90 with a wide-angle lens at most sites, and a Sony RX100V with a Nauticam WWL-1 wide angle lens at Julian Rock Nursery, Cook Island and Flinders Reef. Since the field of view of the two cameras varied, images from the Sony RX100V were batch processed and cropped in ImageJ (Schindelin *et al*., 2012) to ensure comparability, such that each frame captured approximately 1 m^2^ of seabed.

**Figure 1.**
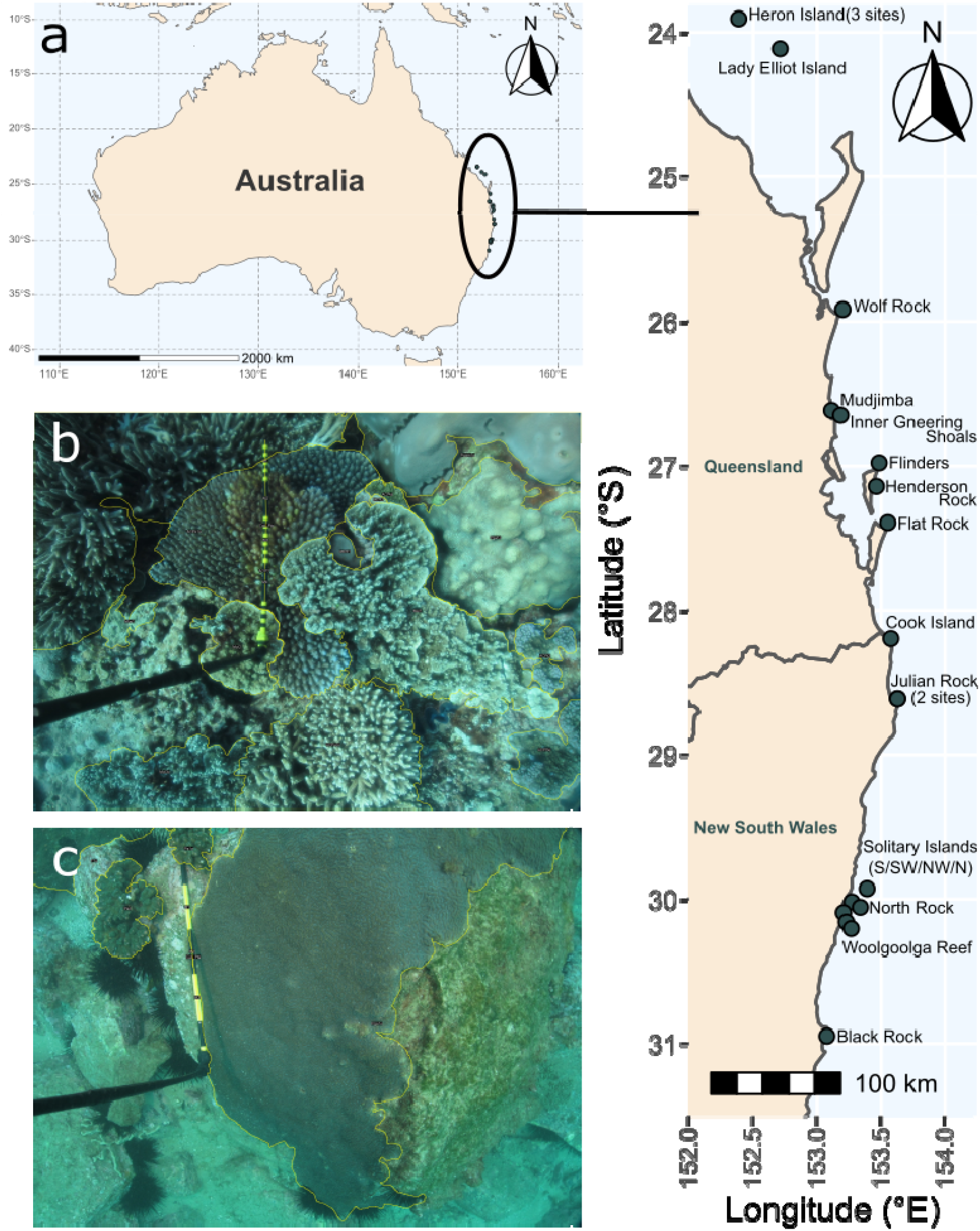
Survey design of the study, showing a) the location of the 20 sampling sites in eastern Australia; image examples of the outlined coral communities from b) Lady Elliot Island and c) Black Rock. The 0.5 m black and yellow graduated calibration stick is visible.

On each image, coral species were visually identified to the lowest taxonomic classification possible (usually genus) using Coral Finder 2021 (Kelley, 2021) and Corals of the World (Veron *et al*., 2016). Coral morphological types were also included and standardised following the classification of Sommer *et al*. (2021). Where variable growth forms are observed for the genera *Montipora, Porites* and *Turbinaria*, they were placed into categories of ‘branching,’ ‘encrusting, ‘laminar’ and ‘massive.’ For *Acropora*, the categories were ‘arborescent,’ ‘corymbose,’ ‘digitate,’ ‘hispidose’ and ‘tabular,’ following Kelley (2021). For each coral colony, the following were recorded: 2D planar area, taxonomic identity, and whether the colony was partially out of frame. This procedure was conducted using the freely available ‘SizeExtractR’ (Lachs *et al*., 2022) workflow in ImageJ (Schindelin *et al*., 2012) and R (R Core Team, 2021). We traced each coral colony manually, added relevant alphanumeric annotations, and compiled the resulting size data into a single database (Lachs *et al*., 2022). In total, 16,598 coral colonies were examined across 1,426 images, capturing 41 coral taxonomic entities (species, genera, family, or groups with uniquely identifiable morphological characteristics; see supporting information for details).

Light limitation, temperature minima, and fluctuations determine the distribution and abundance of corals in our study region (Sommer *et al*., 2018). To characterise and compare long-term environmental trends among our study sites, we extracted 4 km monthly chla (chlorophyll *a* concentration – a proxy for productivity), kd490 (diffuse attenuation coefficient at 490 nm – a proxy for turbidity), and PAR (photosynthetically available radiation) from January 2003 to April 2019 (NOAA, 2022d, 2022b, 2022a); and 1 km monthly Sea Surface Temperature (SST) from June 2002 to May 2019 (NOAA, 2022c). The minima, maxima, means, and standard deviations of each environmental variable were calculated for each site, resulting in a total of 16 variables. A principal component analysis (PCA) was used for dimension reduction of these environmental factors (Figure 2). The first axis (PC1) explains 63% of the observed variance and reflects a gradient from warmer, brighter environments with low turbidity and productivity (negative PC1 scores) to darker, colder environments with high turbidity and productivity (positive PC1 scores). The second axis (PC2), explaining 17% of the variance, is driven by minimum productivity, turbidity, and light availability variability. Negative PC2 scores reflect environments that have the lowest productivity and turbidity, yet unstable light regimes, while positive scores reflect sites whose lowest turbidity and productivity is the least extreme and have the most stable light regimes.

**Figure 2.**
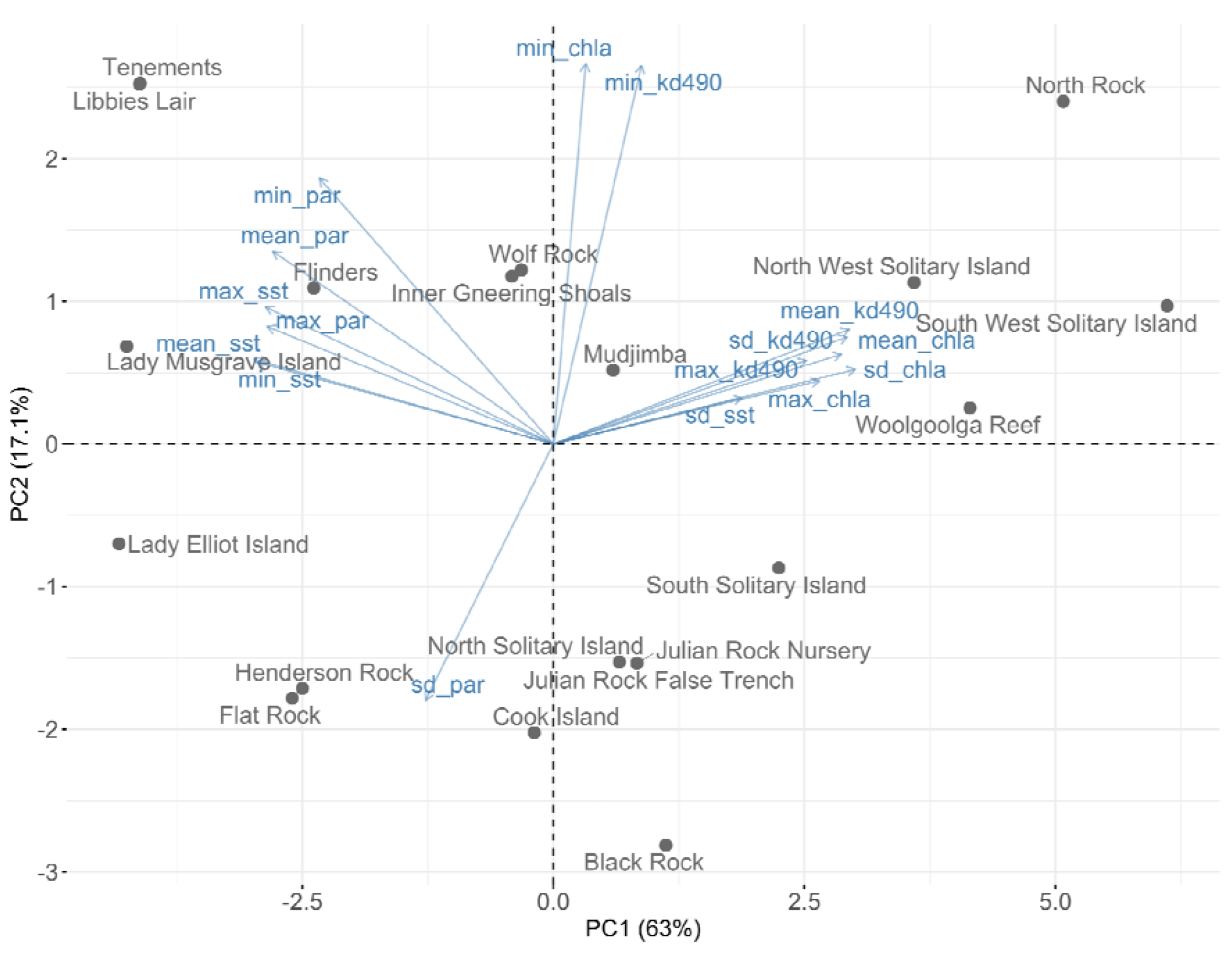
Biplot showing the PCA ordination of our 20 coral populations (Figure 1a) using the 16 environmental variables. Reef names are labelled in grey, the blue arrows are the environmental factors which include the minima (min), maxima (max), means and standard deviations (sd) of chlorophyll *a* concentration (chla), diffuse attenuation coefficient at 490 nm (kd490), sea surface temperature (sst) and photosynthetically available radiation (PAR). The first and second axes jointly explain 80% of the environmental variation in this region.

### Data analyses

Colony sizes were natural log-transformed to normalise their distribution for subsequent analyses and increase the resolution of the highly abundant smaller size classes (Bak & Meesters, 1998). Throughout, log refers to natural logarithm. Colonies marked partially out of frame were excluded as we lacked their true size. This filter resulted in 12,226 coral colonies from 1,322 images, corresponding to 41 coral taxonomic entities. We used two methods to characterise the coral population size structure and establish its relationship with environmental covariates. The first was the calculation of summary statistics (Bak & Meesters, 1998; Adjeroud *et al*., 2007; Anderson & Pratchett, 2014) followed by linear regressions with the scores of PC1 and PC2 and their interaction as explanatory variables. The model combinations were evaluated using Akaike’s Information Criterion (AIC). For each site, the summary statistics calculated were: 1) average coral size (both mean and median), a surrogate for coral age and fecundity (Soong & Lang, 1992). We used the median in the linear regressions as it is not strongly influenced by very small or large colonies, common in our study populations. 2) Coefficient of variation, which allows the comparison of size variation across different sites. 3) Skewness, which measures the asymmetry of size-frequency distributions, with left or right skew indicating the dominance of larger and smaller corals, respectively. 4) Kurtosis, which measures the relative peakedness of a distribution, and has been used to represent growth and recruitment rates (Bak & Meesters, 1998; Adjeroud *et al*., 2007; Anderson & Pratchett, 2014).

We then used compositional functional regressions (Talská *et al*., 2018) to test the effect of environmental covariates (PC1 and PC2 scores) on the entire size-frequency distribution. The benefit of this approach is that the response variable and parameters can be modelled as continuous probability density functions, rather than as simple summary statistics or numbers. Although the general idea of functional regression (the case where the response variables are functions instead of numbers) is known in ecology (*e*.*g*., Yen *et al*., 2015), the standard approach does not ensure that the predicted response function is non-negative and integrates to one (a characteristic of probability density functions, as in our case, size-frequency distributions). This problem can be solved by working in a real vector space (Bayes space), whose elements are continuous probability density functions (Egozcue *et al*., 2006; van den Boogaart *et al*., 2014). Let *f* _1_, *f*_2_ be continuous probability density functions whose support is a closed interval *I* = [a, b] ⊂ ℝ, and let x ∈ ℝ. Then the addition operation in Bayes space is perturbation *f* _1_ ⊕ *f* _2_, defined by

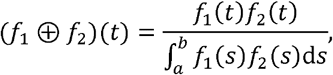

for *t* ∈ *i*, and the scalar multiplication operation is powering *x* ⊙ *f* _1_, defined by

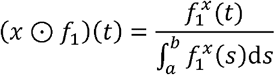

for *t* ∈ *i* (Talská *et al*., 2018). These operations can be thought of as infinite-dimensional versions of the perturbation and powering operations for compositional data, which have been used previously in analyses of the effects of environmental disturbances on coral reef composition and stability (*e*.*g*., Gross & Edmunds, 2015; Vercelloni *et al*., 2020).

Consider the standard linear regression *response = intercept + explanatory variable × coefficient + error*; then the analogous compositional functional regression equation takes the form

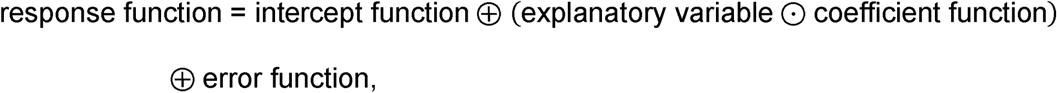

where the error function has a mean of zero. In our particular case, the regression model is

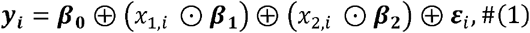

where ***y***_***i***_ is the *response*, a probability density function representing the log coral size-frequency distribution at the *i* th site, the *explanatory variables x*_1,*i*_ and *x*_2,*i*_ are the PC1 and PC2 scores at the *i* th site, the *intercept* ***β***_**0**_ is the size-frequency distribution when each explanatory variable has the value 0, *coefficients* ***β***_**1**_ *and* ***β***_**2**_ are probability density functions describing the effect of a unit increase in PC1 and PC2 respectively on the size-frequency distribution, and the *error* ***ε*** _***i***_ is a probability density function representing the residual or error at the *i* th site.

To obtain estimated continuous size-frequency distributions to use as the response variable, we binned and smoothed the log coral area data from each site. Coral log areas were binned into histograms over the entire observed range across all sites (Talská *et al*., 2018). The number of bins for each site was chosen using Sturges’ rule (Sturges, 1926). Where there were empty bins, we replaced the zeros by 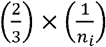, where *n*_*i*_ is the number of corals observed at that site (Martín-Fernández *et al*., 2003; Machalová *et al*., 2021, p. 1053). Then, the size-frequency distributions were centred log-ratio (clr) transformed to give standard addition and scalar multiplication operations, which allows for easier computation (van den Boogaart *et al*., 2014). The clr transformed size-frequency distributions were smoothed using cubic compositional splines (ZB spline basis functions (Machalová *et al*., 2021)) with four knots. The optimum smoothing parameter alpha was chosen by generalized cross validation for each site. The compositional regression model given in Equation 1 was fitted to the binned and smoothed size-frequency distributions (Machalová *et al*., 2021). Approximate 95% confidence bands were obtained as asymptotic normal approximations based on the covariance matrix of estimated coefficients. We calculated pointwise and global R^2^ which measure proportions of variation explained by the model in an analogous way to the usual coefficient of determination (Talská *et al*., 2018).

To determine whether the estimated effects of PC1 and PC2 could be distinguished from zero (no effect), pointwise and global permutation *F*-tests were performed with the observed pointwise *F*-statistic, and its maximum over the whole interval, respectively (Ramsay *et al*., 2009, p. 168). The *F*-tests were carried out by permuting rows of of the ZB-spline coefficients and re-estimating the regression model 10,000 times. We compared observed pointwise and max *F*-statistics with 0.95-quantiles of these statistics from permutations. The residual functions were plotted (and coloured by PC1 score) to check for systematic departures from the model. The coefficient functions ***β***_0_, ***β***_1_ and ***β***_2_ on the clr scale were plotted to visualize the size-frequency distribution at the mean of PC1 and PC2 (***β***_0_) and the effects of each. On the clr scale, positive values of the coefficient functions ***β***_1_ and ***β***_2_ suggest an increase in density at a given log area per unit increase in the explanatory variable, and *vice versa*. Because PC1 seemed to capture most of the environmental variability in our study region, we visualised its effect by plotting the predicted coral size-frequency distributions at the mean value (0) of PC2, for ten equally spaced values of PC1 from its minimum to its maximum.

### Size-biased sampling

Size-frequency distributions estimated from photographs are subject to sampling bias. The larger a coral colony, the less likely it is to fit entirely in the sampling window. Thus, including only those colonies that fit in the sampling window (“minus sampling” (Baddeley, 1998, p. 40)) as we have in this study, biases the estimated size-frequency distribution towards smaller colonies. There are ways to avoid such sampling bias but these require information from outside the sampling window (Baddeley, 1998; sections 2.2-2.4, 2.6; Zvuloni *et al*., 2008), which is unavailable in our data. In the supporting information (S2: Size-biased sampling), we show that this sampling bias does not affect estimates of the coefficient functions for the effects of explanatory variables (***β***_1_ *and* ***β***_2_) in a compositional functional regression, although the bias does affect the estimated intercept function ***β***_0_. Summary statistics and the effects of explanatory variables on the summary statistics will also be subject to sampling bias, but we currently do not have simple solutions to account for these biases.

## Results

### Summary statistics and linear regression

The site with the most coral colonies was Lady Musgrave Island (2,101) while the one with the least was Woolgoolga Reef (38). The median number of corals along this gradient was 526.5 (Q1-Q3: 147.5-718). Statistical summaries of the coral size-frequency distributions are reported in Table S2. Multiple linear regression predicted that the number of coral colonies significantly increased with PC1, but decreased with PC2 (F_2,17_ = 6.80, *P* = 0.007, R^2^ = 0.380) (Figure 3a; Table S3). Thus, more coral colonies were found in warmer, brighter reefs with less turbidity and productivity. Fewer coral colonies were found where light availability was highly variable (more negative PC2 scores) (Figure 3a, blue line). Linear regression showed a positive relationship between PC1 and median log coral colony size (F_1,18_ = 10.7, *P* = 0.004, R^2^ = 0.338), so coral colonies were smaller in warmer, brighter, less productive and less turbid environments and larger in colder, darker, more productive and more turbid environments (Figure 3b; Table S4). In addition, more marginal reef populations such as South West Solitary Island, North Rock and Woolgoolga Reef were more left (negatively) skewed (Figure 4; Table S5; Figure S1) (F_2,17_ = 7.60, *P* = 0.004, R^2^ = 0.410). CV and kurtosis decreased with more positive PC1 scores, suggesting that colony size variation decreased and that the size-frequency distribution became flatter as reefs become more marginal. However, the best models for CV and kurtosis only accounted for a relatively small amount of the variability in our observations (R^2^ = 0.070 and 0.103 respectively; P > 0.05; Table S6-7; Figure S2-3).

**Figure 3.**
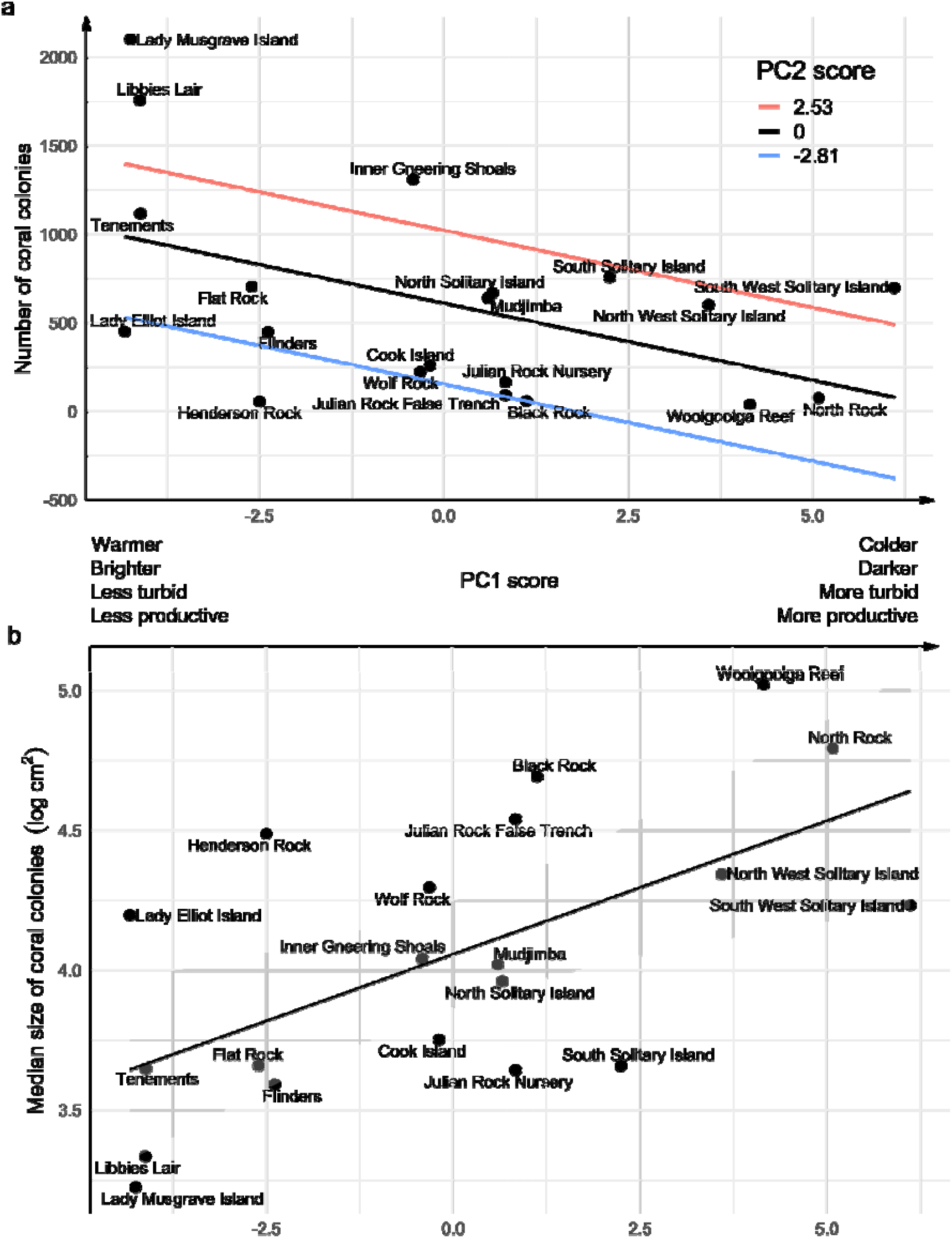
a) The number of coral colonies decreases with PC1. Lines are predictions for the number of coral colonies evaluated at three PC2 scores: the maximum (red), mean (black) and minimum (blue) PC2 scores. b) Median coral colony size increases with PC1. Black line is the line of best fit, and the grey region is the 95% confidence band. The explanatory variables plotted here are chosen based on model selection (Table S3-4). For both a) and b) more positive PC1 scores represent lower sea surface temperature and photosynthetically available radiation (PAR), *i*.*e*., colder and darker, and high chlorophyll *a* concentration and turbidity (kd490), *i*.*e*., more productive and more turbid. In a) more positive PC2 scores represent higher minima of chlorophyll *a* concentration and kd490, but lower standard deviations of PAR.

**Figure 4.**
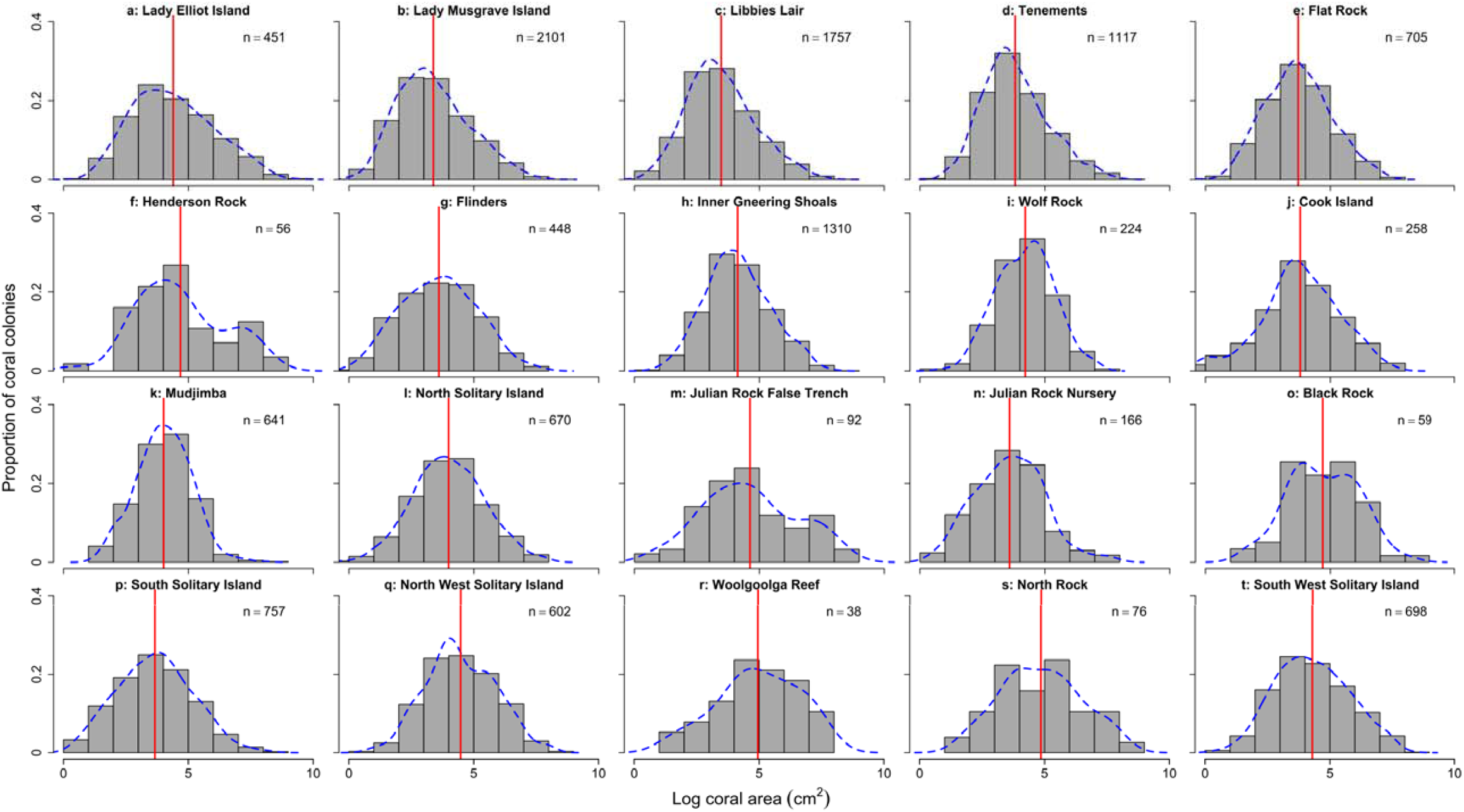
Histograms showing coral colony size structure for each of the 20 reefs. All plots are on the same scale. Blue dashed lines are density estimates. Red lines are mean log coral colony size. Panels (a-t) are ordered from low to high PC1 scores. Increases in PC1 represents increasingly marginal conditions (colder, darker, more turbid and productive waters).

### Compositional functional regression

The smoothed centred log-ratio (clr) densities (Figure S4; analogous to Figure 4) were generally a good representation of our data, except for sites with the lowest number of colonies (*e*.*g*., Black Rock, Henderson Rock, North Rock and Wolf Rock), where estimated densities were artificially high at the lowest log areas.

Compositional functional regressions showed that as PC1 increased, reflecting the transition from warmer, brighter environments to more productive and turbid environments, the mode of the predicted distribution of log coral area moved to the right, and the predicted distribution became broader and flatter (Figure 5, red to blue lines). At the lowest PC1 score, the predicted modal log coral area was approximately 3.5 log cm^2^ (33.1 cm^2^, Figure 5, red), while at the highest PC1 score, the predicted modal log coral area was approximately 5 log cm^2^ (148 cm^2^, Figure 5, blue). Thus, large changes in coral size-frequency distributions along the environmental gradient were plausible. We further showed that increases in PC1 may be associated with lower densities of small to moderate sized corals (∼2-4 log cm^2^) but a higher density of large corals (∼6-7 log cm^2^) (Figure 6, intervals where the 95% confidence band did not cross zero). It was plausible that PC2 had no effect on coral sizes, as the 95% confidence band included zero almost everywhere (Figure S6). The predicted distribution of log area at the mean values of PC1 and PC2 (given by the intercept function ***β***_0_) was symmetrical, with a mode of approximately 4 log cm^2^, corresponding to 54.6 cm^2^ (Figure S5).

**Figure 5.**
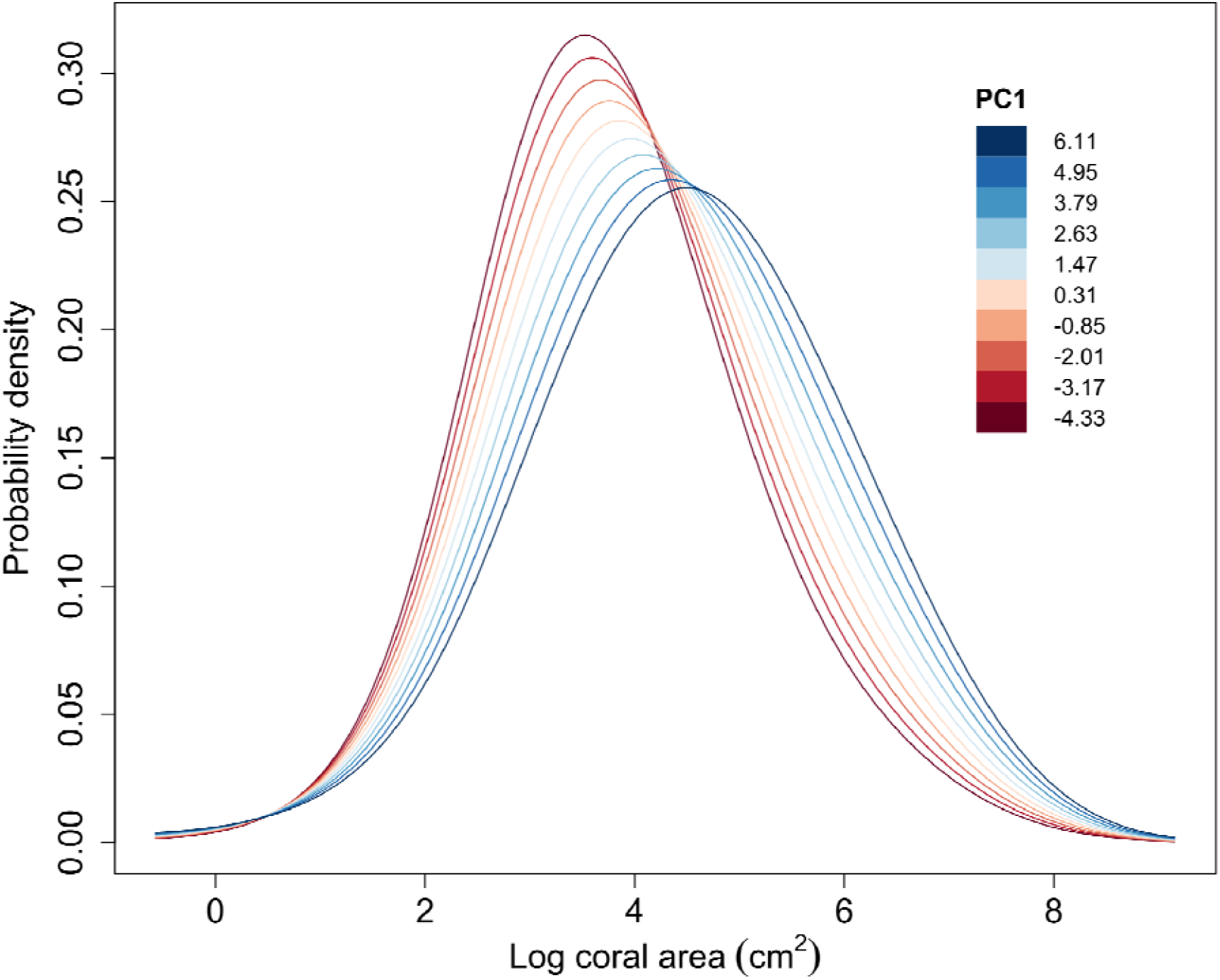
As PC1 increases, the predicted distributions of log coral area become broader and flatter, and the mode increases from ∼3.5 to 5 log cm^2^. Increases in PC1 represents increasingly marginal conditions (colder, darker, more turbid and productive waters). Red to blue lines correspond to predicted distributions for ten equally spaced PC1 scores, from the minimum (−4.33, darkest red) to the maximum (6.11, darkest blue). PC2 values are kept constant at 0 (the mean). The coefficient function determines how the shape of the distribution changes with PC1 but individual distributions are also affected by (the intercept) and thus by the sampling bias (S2: Size-biased sampling).

**Figure 6.**
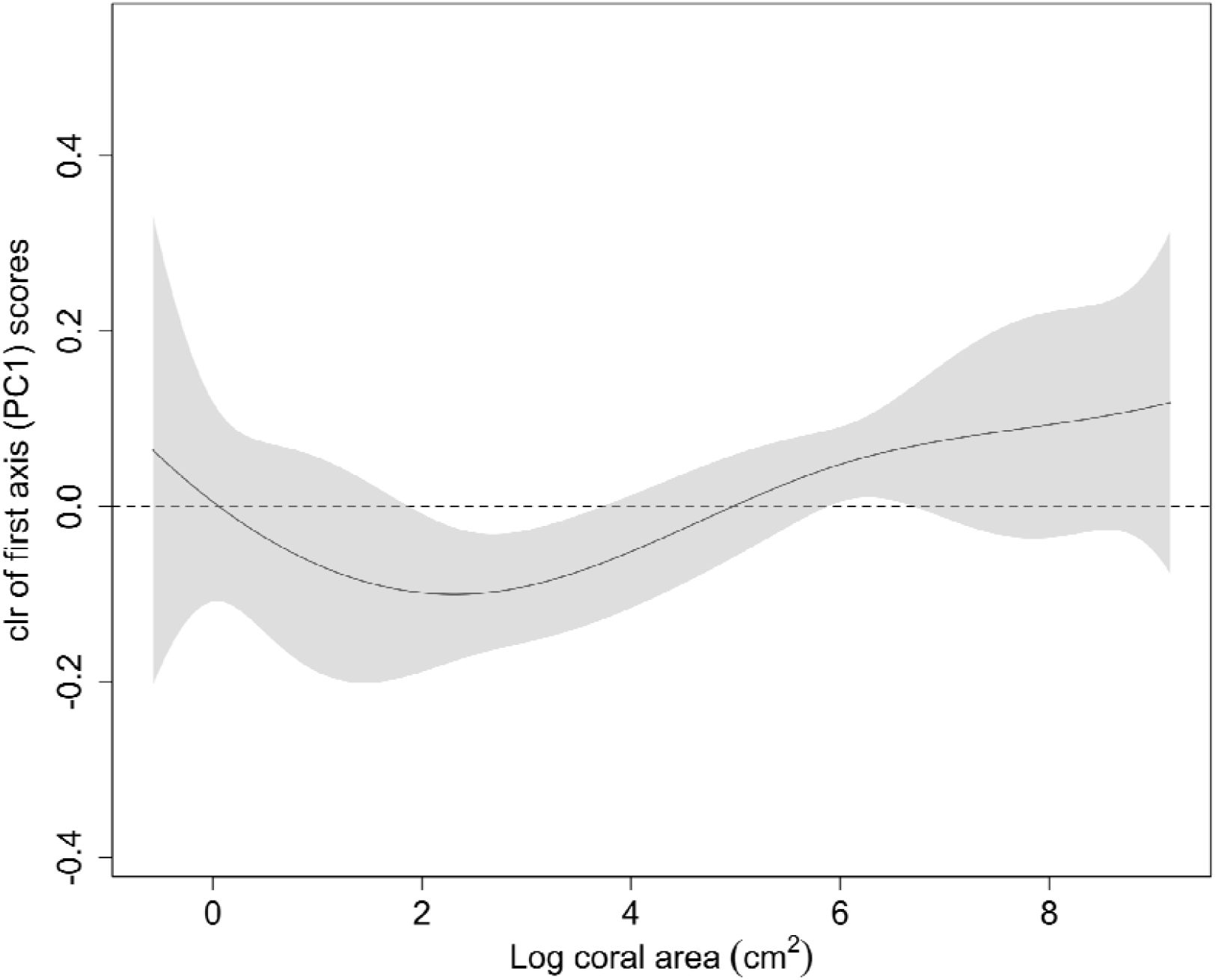
Increases in first axis (PC1) scores mean lower densities of corals at ∼2-4 log cm^2^ but a higher density of corals at ∼6-7 log cm^2^. Increases in PC1 represents increasingly marginal conditions (colder, darker, more turbid and productive waters). The black line is the estimated centred log-ratio (clr) transformation of the coefficient function, which measures the effect of a unit increase in PC1 on the probability density of a given log coral area. Positive values on the y-axis suggest that the corresponding log coral area on the x-axis becomes more likely as PC1 increases, and negative values suggest that the corresponding log coral area becomes less likely. The shaded region is the asymptotic 95% confidence band. The horizontal dashed line represents no effect of PC1 on the probability density of log coral area.

The global R^2^ for our model was 0.19, so that the model explained relatively little of the variation in size-frequency distributions, although with higher amounts of variation explained at log coral sizes between 3 and 6 (Figure S7). Similar peaks were observed for the pointwise *F* test statistics (Figure S8). However, because the maximum pointwise *F* statistic (Figure S8, dotted line) did not exceed the 0.95-quantile of the distribution of such maxima anywhere (Figure S8, dashed line), it was plausible that neither PC1 nor PC2 affected coral size-frequency distributions.

Residual plots showed little systematic pattern in the residual functions, except that they tended to be further from zero at the smallest or largest coral areas (Figure S9). This might be associated with smoothing artefacts (Machalová *et al*., 2021), where there are lower numbers of colonies at the extreme sizes.

## Discussion

Understanding the drivers of change in population size structure is fundamental to robust predictions of population dynamics (Edmunds & Riegl, 2020; Edmunds, 2021). Here, examining 20 reefs along the tropical to subtropical transition zone in Eastern Australia, we found fewer but bigger corals in sites characterised by greater environmental stress and temporal variability compared to sites that have a more stable environmental regime. Despite some uncertainty in compositional functional regression results, it is plausible that the high coral cover in Australian high-latitude coral communities (Harriott *et al*., 1994) is created by few large coral colonies. The lower growth rates and higher fission rates of larger corals (Dornelas *et al*., 2017) could be the main driver of coral persistence in marginal reefs (Cant *et al*., 2022). As ongoing climate change leads to more variable and extreme environmental conditions (Spady *et al*., 2022), we hypothesise that tropical accreting reefs will increasingly exhibit a population size structure that resembles those now observed at marginal reefs; namely, one with a higher proportion of larger corals. Our research suggests that future reef persistence might be governed by low growth and recruitment, and be reliant on the survival of larger corals (Bak & Meesters, 1999; Cant *et al*., 2020; Dietzel *et al*., 2020).

To our knowledge, this is the first study to use compositional functional regression (Talská *et al*., 2018) to examine population size structure changes along a large biogeographic gradient, and we demonstrate its potential. The ability to account for the entire probability density curve is elegant, as one can examine with higher resolution how and which coral sizes are affected by the environmental drivers. Compositional functional regression also removes the need to run multiple linear regressions on arbitrarily chosen summary statistics (Talská *et al*., 2018), which can be difficult to interpret. In principle, it is also possible to calculate how proportional population growth rate depends on an explanatory variable (*e*.*g*. PC1 in our case), making this method relevant for demography. A compositional functional regression with one explanatory variable is a map from the real line (the explanatory variable) to Bayes space (size-frequency distributions) (Egozcue *et al*., 2013, section 4), while a stationary size-structured integral projection model can be thought of as a map from Bayes space (size-frequency distributions) to the real line (stationary proportional population growth rates) (Easterling *et al*., 2000). These properties mean that the derivative of the proportional population growth rate with respect to the environmental variable can be calculated using the chain rule. In contrast, we do not have maps from summary statistics such as skewness or kurtosis to proportional population growth rate, because these summaries do not contain all the information needed to calculate proportional population growth rate. Thus, summary statistics are dead ends in quantitative terms.

Although both linear regression and compositional functional regression support the main finding of fewer but bigger corals in marginal reefs, the evidence in the latter is weaker. This difference could simply be methodological, *i*.*e*., having to consider the effect of environmental covariates over the entire size-frequency distribution, instead of a single value (of a summary statistic) for each reef. It could also be due to higher sampling variability (noise) in the fitting of histograms and centred log-ratio densities at sites where there were fewer corals, and this variability is not accounted for in our current approach. Future work on population size structure could sample from a larger area on marginal reefs, where there are fewer coral colonies to gather more coral size data; or use sampling methods that can consistently capture the size of whole corals, so as to reduce the number of colonies not included in the analyses. Other factors that were not captured by our analyses could also have acted on the coral size-frequency distributions at each reef, *e*.*g*., wave exposure and reef topography. Corals of different sizes and growth forms have been shown to have differential resistance of being overturned by storm waves (Madin *et al*., 2014).

Reefs with higher rugosity and thus complexity have been shown to support more smaller corals (Crabbe, 2010). Competition can also affect coral sizes, such as competition for space with other non-coral, sessile benthic organisms like algae, corallimorpharians and zoanthids (Abrego *et al*., 2021; Reimer *et al*., 2021). These biotic interactions may reduce the rate at which corals grow (Chadwick & Morrow, 2011), so that reefs with a higher cover of other benthic taxa could have fewer or smaller corals than expected. Finally, although the environmental gradient appears to select for larger corals, the taxonomic identity and morphology of the coral also determine their life history (Darling *et al*., 2012). For example, the encrusting *Micromussa lordhowenesis*, which is commonly observed on subtropical reefs in this region, is generally much smaller than the laminar *Tubinaria*. Where taxa are abundant and sample sizes large enough, it will be meaningful to investigate specific responses to forecast population dynamics under global change.

Our work assessing coral population size structure over a large biogeographic scale offers a glimpse into a possible response of coral assemblages to climate change. Our main finding of increasingly marginal conditions selecting for fewer but larger coral colonies, echoes previous findings that larger corals remain post-disturbance (Bak & Meesters, 1998; Dietzel *et al*., 2020; Lachs *et al*., 2021), but see also Pisapia *et al*. (2020) for examples of colonies becoming smaller. The ecological mechanisms that can lead to the prevalence of bigger corals are many, complex and interactive, including a combination of low recruitment, partial mortality and slow growth. We hypothesise that the effects of ongoing climate change, which lead to more marginal conditions, could select mechanisms that shift reef coral population size structure towards a larger proportion of bigger individuals. Such a shift is concerning because coral populations with more smaller corals (juveniles) are usually considered to have a higher recovery potential after disturbance events (Riegl *et al*., 2012; Pisapia *et al*., 2019; Dietzel *et al*., 2020; Lachs *et al*., 2021). Smaller corals thus have a disproportional importance for population persistence, and may be a useful target for conservation. As such, we recommend improving our understanding of coral reproduction and recruitment dynamics along latitudinal gradients (*e*.*g*., Mizerek *et al*., 2021), as it can provide an insight into how reef populations persist and recover despite suboptimal conditions. In addition to understanding larval dispersal and ensuring population connectivity (Greiner *et al*., 2022), there is a continual need to explore other mechanisms that facilitate rapid reef recovery. For example, Doropoulos *et al*. (2022) hypothesised that coral cryptic seed banks play an important role in reef recovery after cyclone disturbances.

Climate change is projected to continue to affect population dynamics worldwide (Lawson *et al*., 2015). Thus, it remains pertinent for ecologists to examine changes in population size structure at biogeographic scales through time (*e*.*g*., Riegl *et al*., 2012; Dietzel *et al*., 2021). Advances in compositional functional regression (Talská *et al*., 2018) provide a comprehensive demographic tool for ecologists to examine population size structure, allowing us to gain insight into how environmental extremes and variabilities affect population dynamics (Kreyling *et al*., 2014). Collectively, our work on the coral population size structure of reefs in the Eastern Australian biogeographic transition zone highlights fundamental differences along the ∼ 900 km tropical to subtropical gradient, where bigger corals are likely selected for in marginal conditions. Marginal reefs and future coral reefs could, therefore, become less resilient to disturbances like marine heatwaves due to a lowered recovery potential.

## Notes

### Competing Interest Statement

The authors have declared no competing interest.

