## Supporting information for "High-latitude marginal reefs support fewer but bigger corals than their tropical counterparts"

### S1: Supplemental tables and figures

Table S1. GPS coordinates and the sampling date of the 20 sites, ordered by decreasing latitude.

|  | Site name | Latitude (°) | Longitude (°) | Sampling date (dd/mm/yyyy) |
| --- | --- | --- | --- | --- |
| 1. | Black Rock | -30.94837 | 153.0761 | 10/09/2018 |
| 2. | South Solitary Island | -30.20478 | 153.2652 | 13/09/2018 |
| 3. | South West Solitary Island | -30.15921 | 153.2281 | 09/09/2018 |
| 4. | Woolgoolga Reef | -30.09374 | 153.2056 | 19/10/2016 |
| 5. | North West Solitary Island | -30.01897 | 153.2697 | 12/09/2018 |
| 6. | North Rock | -29.97339 | 153.2572 | 16/04/2016 |
| 7. | North Solitary Island | -29.92772 | 153.3896 | 11/09/2018 |
| 8. | Julian Rock False Trench | -28.61257 | 153.6286 | 31/05/2016 |
| 9. | Julian Rock Nursery | -28.61087 | 153.6281 | 14/09/2018 |
| 10. | Cook Island | -28.19627 | 153.5763 | 07/09/2018 |
| 11. | Flat Rock | -27.39306 | 153.5522 | 15/08/2010 |
| 12. | Henderson Rock | -27.13161 | 153.4781 | 18/03/2011 |
| 13. | Flinders Reef | -26.97765 | 153.4841 | 18/09/2018 |
| 14. | Inner Gneering Shoals | -26.64858 | 153.1834 | 29/08/2012 |
| 15. | Mudjimba | -26.61614 | 153.1130 | 28/08/2012 |
| 16. | Wolf Rock | -25.91667 | 153.2000 | 09/08/2010 |
| 17. | Lady Elliot Island | -24.11500 | 152.7095 | 27/09/2018 |
| 18. | Lady Musgrave Island | -23.90603 | 152.3870 | 26/09/2018 |
| 19. | Libbies Lair | -23.43458 | 151.9336 | 23/09/2018 |
| 20. | Tenements | -23.43274 | 151.9293 | 24/09/2018 |

Table S2. Statistical summaries of the size-frequency distribution at the twenty reefs. All areas are in log cm^2^. Numbers are rounded to three decimal places where appropriate. Sites are ordered from the lowest to the highest PC1 scores from top to bottom. Increasing PC1 scores represent lower sea surface temperature and photosynthetically available radiation (PAR), *i.e.,* colder and darker, and high chlorophyll a concentration and turbidity (kd490), *i.e.,* more productive and more turbid.

| Site | Number of coral colonies | Mean coral area | Median coral area | Standard deviation of coral area | Coefficient of variation (CV) | Skewness | Kurtosis |
| --- | --- | --- | --- | --- | --- | --- | --- |
| Lady Elliot Island | 451 | 4.393 | 4.197 | 1.617 | 36.809 | 0.346 | 2.485 |
| Lady Musgrave Island | 2101 | 3.379 | 3.224 | 1.441 | 42.646 | 0.434 | 2.776 |
| Libbies Lair | 1757 | 3.469 | 3.335 | 1.37 | 39.493 | 0.421 | 2.967 |
| Tenements | 1117 | 3.814 | 3.648 | 1.302 | 34.137 | 0.499 | 3.023 |
| Flat Rock | 705 | 3.711 | 3.659 | 1.325 | 35.705 | 0.171 | 2.724 |
| Henderson Rock | 56 | 4.683 | 4.489 | 1.742 | 37.198 | 0.243 | 2.521 |
| Flinders | 448 | 3.599 | 3.592 | 1.524 | 42.345 | 0.076 | 2.521 |
| Inner Gneering Shoals | 1310 | 4.131 | 4.04 | 1.301 | 31.494 | 0.211 | 2.729 |
| Wolf Rock | 224 | 4.216 | 4.296 | 1.147 | 27.206 | -0.016 | 2.721 |
| Cook Island | 258 | 3.794 | 3.752 | 1.546 | 40.749 | -0.273 | 3.094 |
| Mudjimba | 641 | 4.008 | 4.019 | 1.106 | 27.595 | 0.068 | 3.044 |
| North Solitary Island | 670 | 3.988 | 3.961 | 1.436 | 36.008 | 0.057 | 2.897 |
| Julian Rock False Trench | 92 | 4.624 | 4.541 | 1.881 | 40.679 | 0.14 | 2.303 |
| Julian Rock Nursery | 166 | 3.588 | 3.643 | 1.378 | 38.406 | 0.215 | 3.018 |
| Black Rock | 59 | 4.695 | 4.693 | 1.398 | 29.776 | 0.004 | 2.481 |
| South Solitary Island | 757 | 3.661 | 3.658 | 1.511 | 41.273 | 0.15 | 2.696 |
| North West Solitary Island | 602 | 4.475 | 4.343 | 1.41 | 31.508 | 0.102 | 2.647 |
| Woolgoolga Reef | 38 | 4.949 | 5.02 | 1.654 | 33.421 | -0.281 | 2.361 |
| North Rock | 76 | 4.857 | 4.793 | 1.597 | 32.88 | 0.096 | 2.308 |
| South West Solitary Island | 698 | 4.294 | 4.233 | 1.48 | 34.467 | 0.127 | 2.496 |

Table S3. Linear regressions showing the relationship between number of coral colonies and the PC1 and PC2 scores. The best model (lowest AIC) is in bold. PC1 captures the differences in sea surface temperature, photosynthetically available radiation, high chlorophyll a concentration and turbidity (kd490). More positive PC2 scores mean higher minima of chlorophyll a concentration and kd490, but lower standard deviations of PAR. Values unless stated otherwise are corrected to three significant figures.

| Model formula | Predictor | Estimate | Std. Error | *t* value | P value | Adjusted R^2^ | AIC (2 d.p.) |
| --- | --- | --- | --- | --- | --- | --- | --- |
| Coral count ~ PC1 | Intercept | 611 | 116 | 5.24 | < 0.001 | 0.185 | 252.16 |
|  | PC1 | -87.0 | 37.7 | -2.31 | 0.0332 |  |  |
| Coral count ~ PC2 | Intercept | 611 | 118 | 5.20 | < 0.001 | 0.173 | 252.46 |
|  | PC2 | 162 | 72.8 | 2.23 | 0.0388 |  |  |
| **Coral count ~ PC1 + PC2** | Intercept | 611 | 102 | 6.01 | < 0.001 | 0.379 | **247.58** |
|  | PC1 | -87.0 | 32.9 | -2.64 | 0.0171 |  |  |
|  | PC2 | 162 | 63.1 | 2.57 | 0.0198 |  |  |
| Coral count ~ PC1 * PC2 | Intercept | 611 | 101 | 6.05 | < 0.001 | 0.388 | 248.06 |
|  | PC1 | -61.5 | 39.8 | -1.55 | 0.142 |  |  |
|  | PC2 | 151 | 63.4 | 2.39 | 0.0298 |  |  |
|  | PC1:PC2 | -27.8 | 24.8 | -1.12 | 0.278 |  |  |

Table S4. Linear regressions showing the relationship between the median coral colony size and the PC1 and PC2 scores. The best model (lowest AIC) is in bold. PC1 captures the differences in sea surface temperature, photosynthetically available radiation, high chlorophyll a concentration and turbidity (kd490). More positive PC2 scores mean higher minima of chlorophyll a concentration and kd490, but lower standard deviations of PAR. Values are corrected to three significant figures.

| Model formula | Predictor | Estimate | Std. Error | *t* value | P value | Adjusted R^2^ | AIC |
| --- | --- | --- | --- | --- | --- | --- | --- |
| **Median size ~ PC1** | Intercept | 4.06 | 0.0902 | 45.0 | < 0.001 | 0.338 | **-34.4** |
|  | PC1 | 0.0954 | 0.0292 | 3.27 | 0.00423 |  |  |
| Median size ~ PC2 | Intercept | 4.06 | 0.113 | 35.9 | < 0.001 | -0.0366 | -25.4 |
|  | PC2 | -0.0402 | 0.0700 | -0.574 | 0.573 |  |  |
| Median size ~ PC1 + PC2 | Intercept | 4.06 | 0.0915 | 44.3 | < 0.001 | 0.319 | -33.0 |
|  | PC1 | 0.0955 | 0.0296 | 3.23 | 0.00495 |  |  |
|  | PC2 | -0.0402 | 0.0567 | -0.709 | 0.488 |  |  |
| Median size ~ PC1 * PC2 | Intercept | 4.06 | 0.0902 | 45.0 | < 0.001 | 0.339 | -32.8 |
|  | PC1 | 0.0706 | 0.0355 | 1.99 | 0.0642 |  |  |
|  | PC2 | -0.0293 | 0.0566 | -0.518 | 0.611 |  |  |
|  | PC1:PC2 | 0.0272 | 0.0221 | 1.23 | 0.237 |  |  |

Table S5. Linear regressions showing the relationship between skewness and the PC1 and PC2 scores. The best model (lowest AIC) is in bold. PC1 captures the differences in sea surface temperature, photosynthetically available radiation, high chlorophyll a concentration and turbidity (kd490). More positive PC2 scores mean higher minima of chlorophyll a concentration and kd490, but lower standard deviations of PAR. Values are corrected to three significant figures.

| Model formula | Predictor | Estimate | Std. Error | *t* value | P value | Adjusted R^2^ | AIC |
| --- | --- | --- | --- | --- | --- | --- | --- |
| Skewness ~ PC1 | Intercept | 0.140 | 0.0372 | 3.75 | 0.00146 | 0.317 | -69.9 |
|  | PC1 | -0.0377 | 0.0120 | -3.14 | 0.00571 |  |  |
| Skewness ~ PC2 | Intercept | 0.140 | 0.0434 | 3.22 | 0.00481 | 0.0699 | -63.7 |
|  | PC2 | 0.0419 | 0.0269 | 1.56 | 0.137 |  |  |
| **Skewness ~ PC1 + PC2** | Intercept | 0.140 | 0.0345 | 4.04 | < 0.001 | 0.410 | **-71.9** |
|  | PC1 | -0.0376 | 0.0112 | -3.37 | 0.00361 |  |  |
|  | PC2 | 0.0419 | 0.0214 | 1.96 | 0.0670 |  |  |
| Skewness ~ PC1 * PC2 | Intercept | 0.140 | 0.0356 | 3.912 | 0.00123 | 0.374 | -70.0 |
|  | PC1 | -0.0387 | 0.0140 | -2.76 | 0.0139 |  |  |
|  | PC2 | 0.0424 | 0.0223 | 1.90 | 0.0762 |  |  |
|  | PC1:PC2 | 0.00112 | 0.00874 | 0.128 | 0.900 |  |  |

Table S6. Linear regressions showing the relationship between the coefficient of variation and the PC1 and PC2 scores. The best model (lowest AIC) is in bold. PC1 captures the differences in sea surface temperature, photosynthetically available radiation, high chlorophyll a concentration and turbidity (kd490). More positive PC2 scores mean higher minima of chlorophyll a concentration and kd490, but lower standard deviations of PAR. Values are corrected to three significant figures.

| Model formula | Predictor | Estimate | Std. Error | *t* value | P value | Adjusted R^2^ | AIC |
| --- | --- | --- | --- | --- | --- | --- | --- |
| **CV ~ PC1** | Intercept | 35.7 | 1.02 | 35.0 | < 0.001 | 0.0701 | **62.6** |
|  | PC1 | -0.514 | 0.329 | -1.56 | 0.136 |  |  |
| CV ~ PC2 | Intercept | 35.7 | 1.06 | 33.5 | < 0.001 | -0.0149 | 64.3 |
|  | PC2 | -0.561 | 0.660 | -0.85 | 0.407 |  |  |
| CV ~ PC1 + PC2 | Intercept | 35.7 | 1.02 | 34.8 | < 0.001 | 0.0585 | 63.7 |
|  | PC1 | -0.514 | 0.331 | -1.55 | 0.139 |  |  |
|  | PC2 | -0.561 | 0.636 | -0.882 | 0.390 |  |  |
| CV ~ PC1 * PC2 | Intercept | 35.7 | 1.05 | 33.9 | < 0.001 | 0.00795 | 65.5 |
|  | PC1 | -0.428 | 0.414 | -1.03 | 0.317 |  |  |
|  | PC2 | -0.598 | 0.660 | -0.906 | 0.378 |  |  |
|  | PC1:PC2 | -0.0942 | 0.258 | -0.365 | 0.720 |  |  |

Table S7. Linear regressions showing the relationship between kurtosis and the PC1 and PC2 scores. The best model (lowest AIC) is in bold. PC1 captures the differences in sea surface temperature, photosynthetically available radiation, high chlorophyll a concentration and turbidity (kd490). More positive PC2 scores mean higher minima of chlorophyll a concentration and kd490, but lower standard deviations of PAR. Values are corrected to three significant figures.

| Model formula | Predictor | Estimate | Std. Error | *t* value | P value | Adjusted R^2^ | AIC |
| --- | --- | --- | --- | --- | --- | --- | --- |
| **Kurtosis ~ PC1** | Intercept | 2.69 | 0.0538 | 50.0 | < 0.001 | 0.103 | **-55.1** |
|  | PC1 | -0.0309 | 0.0174 | -1.78 | 0.0918 |  |  |
| Kurtosis ~ PC2 | Intercept | 2.69 | 0.0582 | 46.2 | < 0.001 | -0.0535 | -51.9 |
|  | PC2 | 0.00685 | 0.0361 | 0.19 | 0.852 |  |  |
| Kurtosis ~ PC1 + PC2 | Intercept | 2.69 | 0.0553 | 48.7 | < 0.001 | 0.0520 | -53.2 |
|  | PC1 | -0.0309 | 0.0179 | -1.73 | 0.101 |  |  |
|  | PC2 | 0.00685 | 0.0343 | 0.200 | 0.844 |  |  |
| Kurtosis ~ PC1 * PC2 | Intercept | 2.69 | 0.0523 | 51.4 | < 0.001 | 0.151 | -54.6 |
|  | PC1 | -0.0107 | 0.0206 | -0.518 | 0.611 |  |  |
|  | PC2 | -0.00200 | 0.0328 | -0.061 | 0.952 |  |  |
|  | PC1:PC2 | -0.0221 | 0.0128 | -1.73 | 0.104 |  |  |

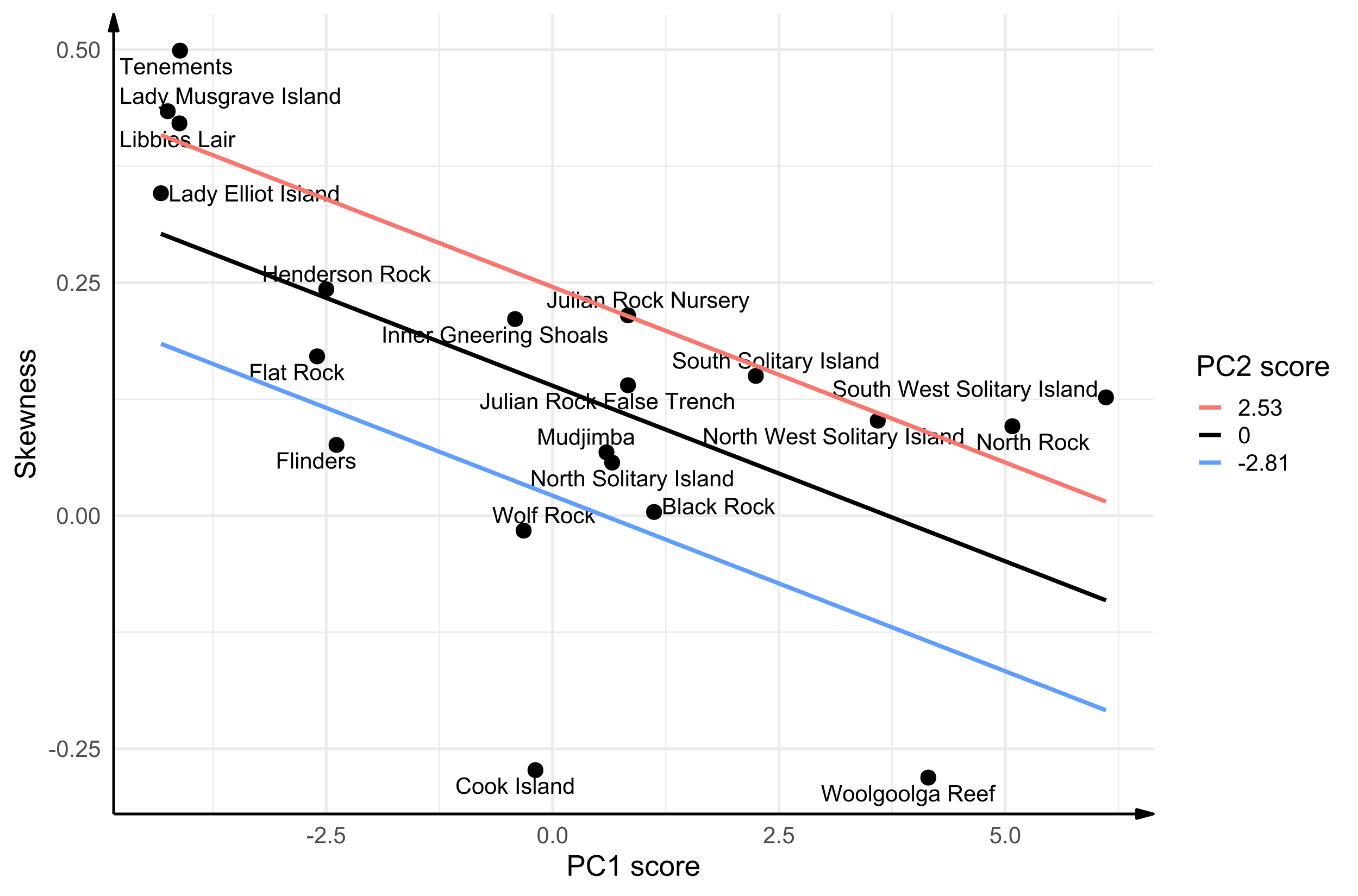

Figure S1. Skewness of the coral size-frequency distribution decreases with increasing PC1 but increases with PC2. Lines are predictions for skewness evaluated at three PC2 scores: the maximum (red), mean (black) and minimum (blue) PC2 scores. More positive PC1 scores represent lower sea surface temperature and photosynthetically available radiation (PAR), *i.e.,* colder and darker, and high chlorophyll a concentration and turbidity (kd490), *i.e.,* more productive and more turbid. More positive PC2 scores represent higher minima of chlorophyll a concentration and kd490, but lower standard deviations of PAR.

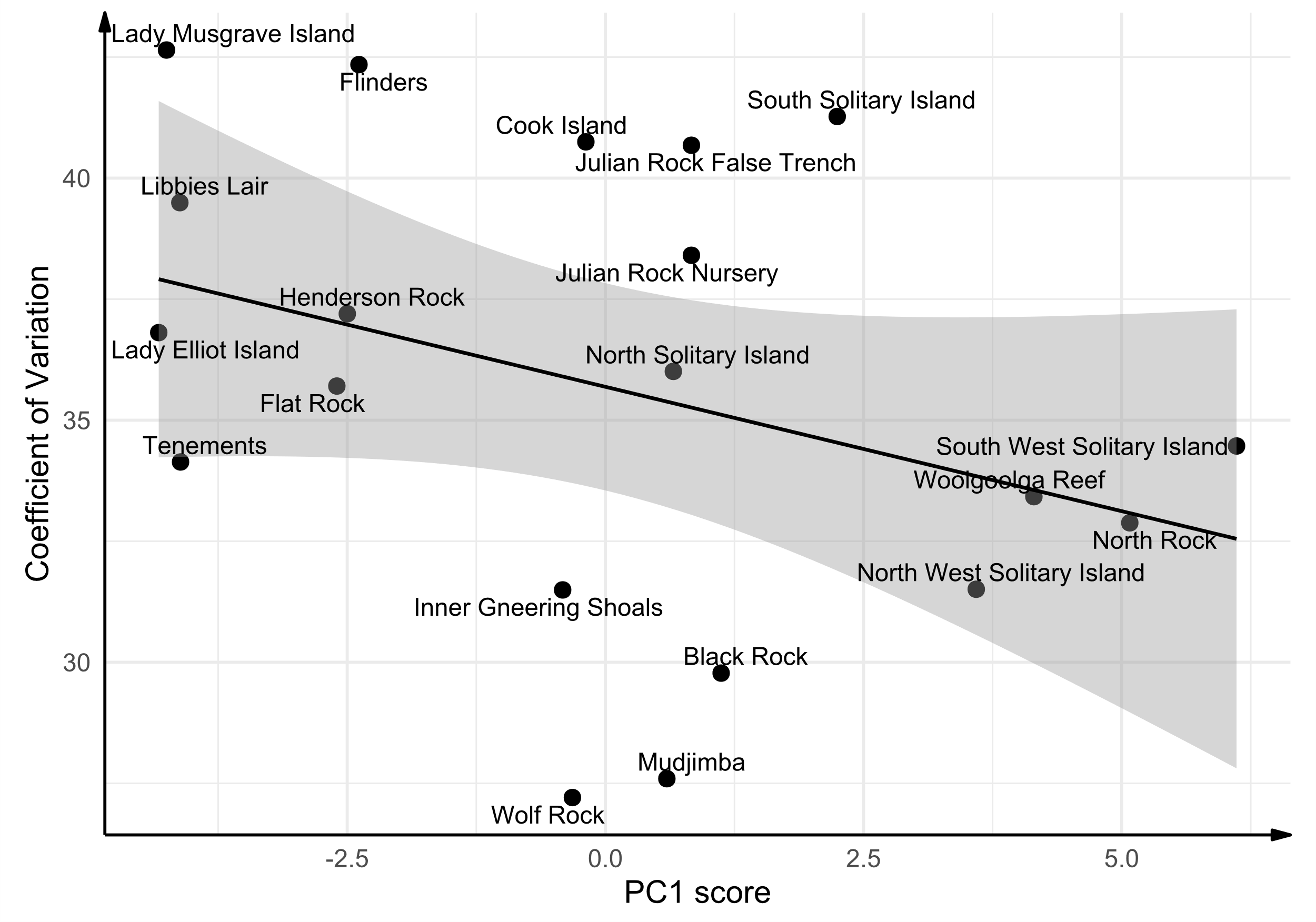

Figure S2. The colony size coefficient of variation (CV) decreases with PC1. PC1 is fitted as the explanatory variable here because of model selection (Table S6). Black line is the line of best fit, and the grey region is the 95% confidence band. More positive PC1 scores represent lower sea surface temperature and photosynthetically available radiation (PAR), *i.e.,* colder and darker, and high chlorophyll *a* concentration and turbidity (kd490), *i.e.,* more productive and more turbid.

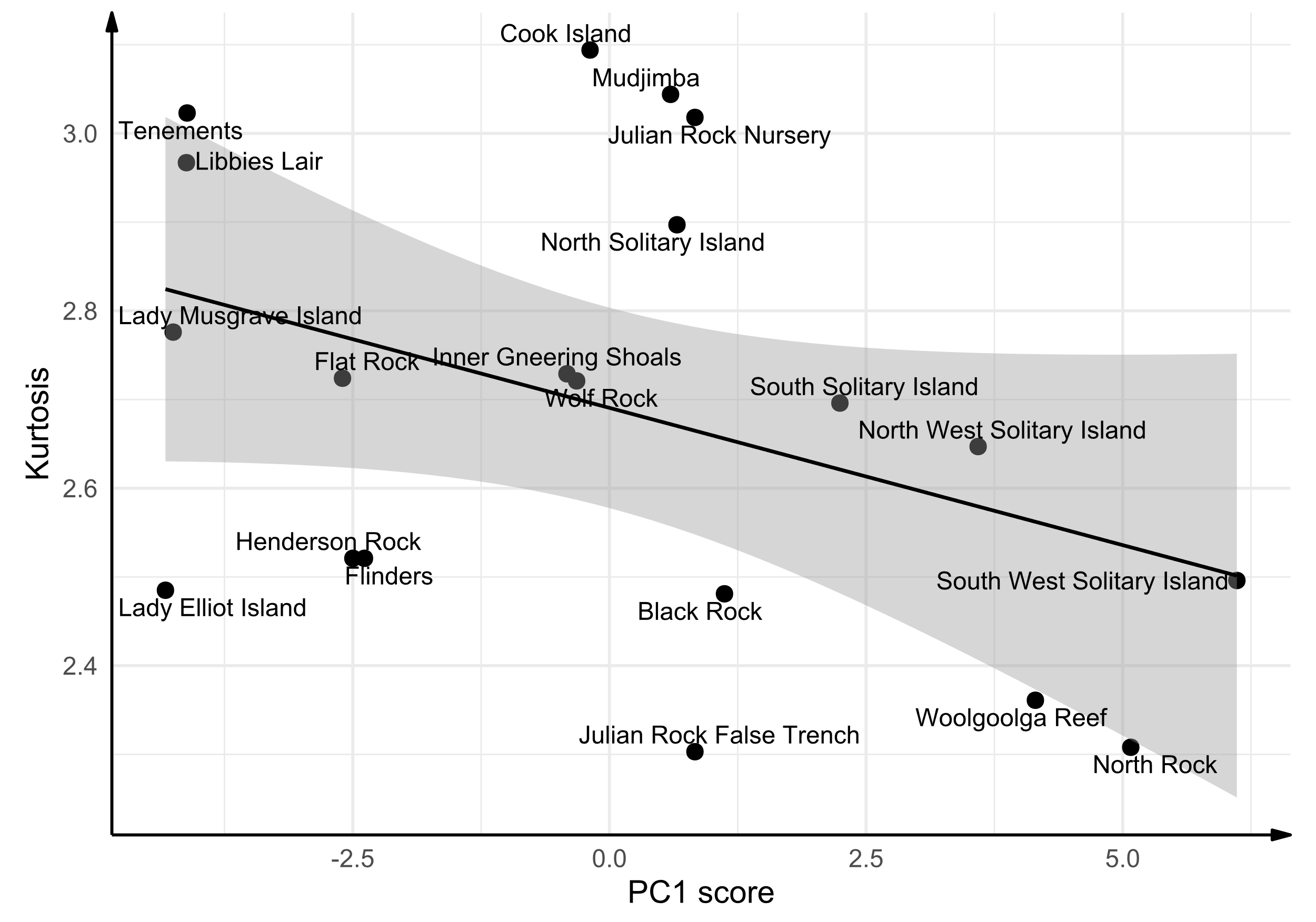

Figure S3. Kurtosis of the coral size-frequency distribution decreases with PC1. PC1 is fitted as the explanatory variable here because of model selection (Table S7). Black line is the line of best fit, and the grey region is the 95% confidence band. More positive PC1 scores represent lower sea surface temperature and photosynthetically available radiation (PAR), *i.e.,* colder and darker, and high chlorophyll a concentration and turbidity (kd490), *i.e.,* more productive and more turbid.

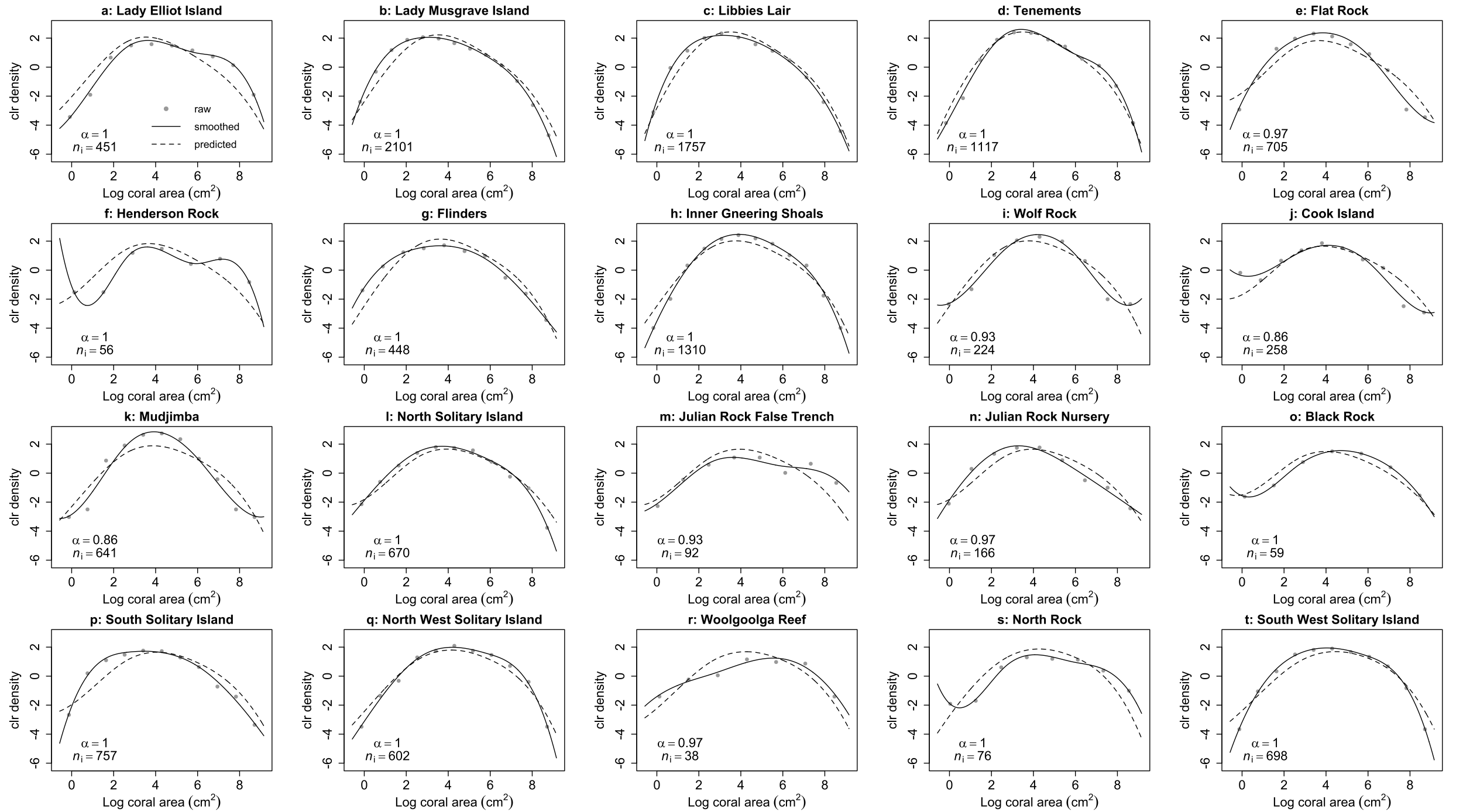

Figure S4. Centred log-ratio (clr) transformed densities of log coral area for the 20 reefs, ordered from low to high PC1 scores (left to right; top to bottom). Grey dots are binned raw data points, black solid lines are smoothed densities and dotted lines are predicted clr densities from a compositional functional regression. For each site, *α* is the value of the smoothing parameter and *n_i_* is the number of corals at the site.

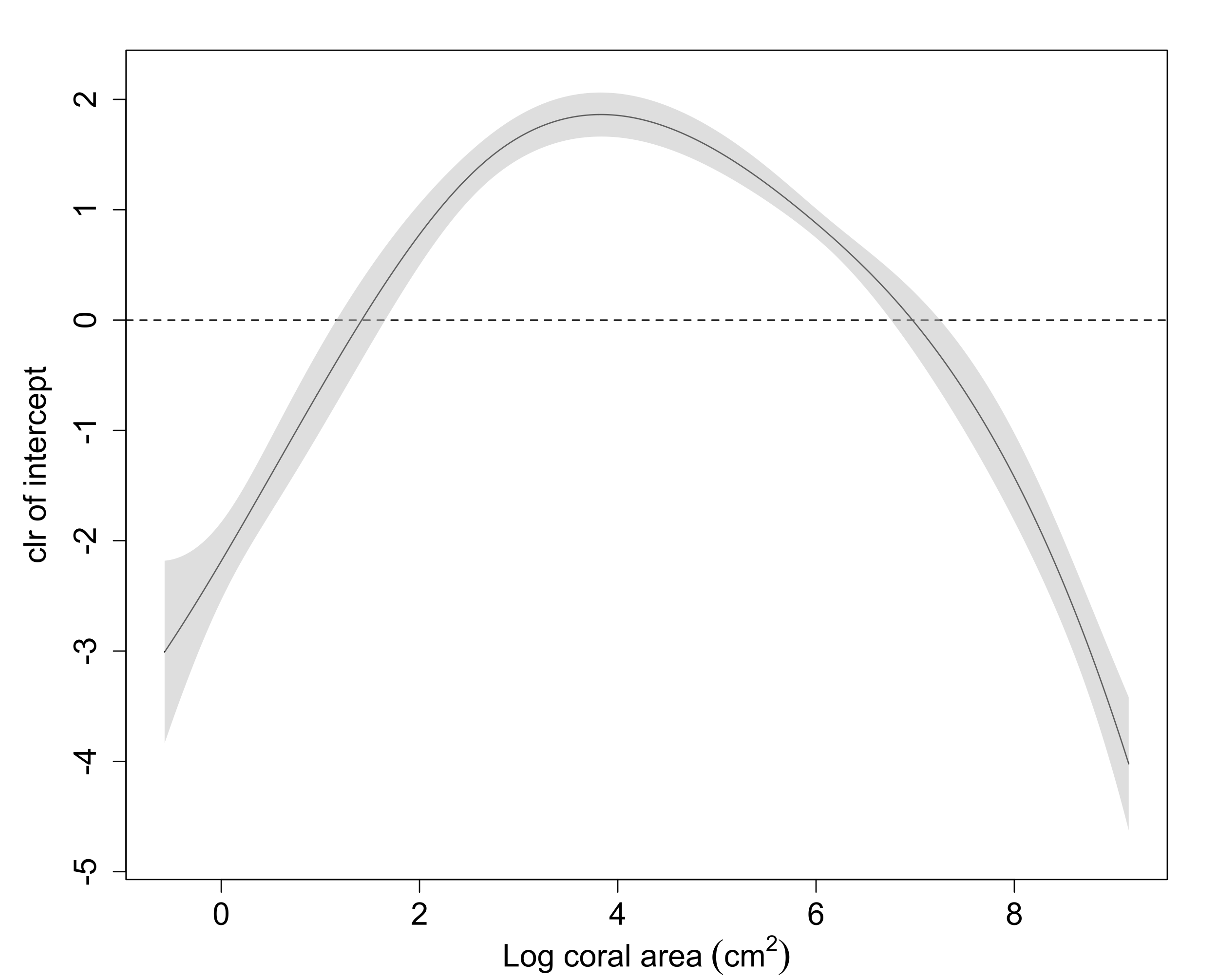

Figure S5. When the mean values of PC1 and PC2 are both zero, the coral size-frequency distribution is symmetrical. The black line is the estimated centred log-ratio (clr) transformation of the coefficient function $\boldsymbol{\beta}_{0}$ (the intercept), which is the prediction when PC1 and PC2 are both equal to zero. The grey shaded region is the asymptotic 95% confidence band. The horizontal dashed line represents no effect of the intercept on the probability density of log coral area.

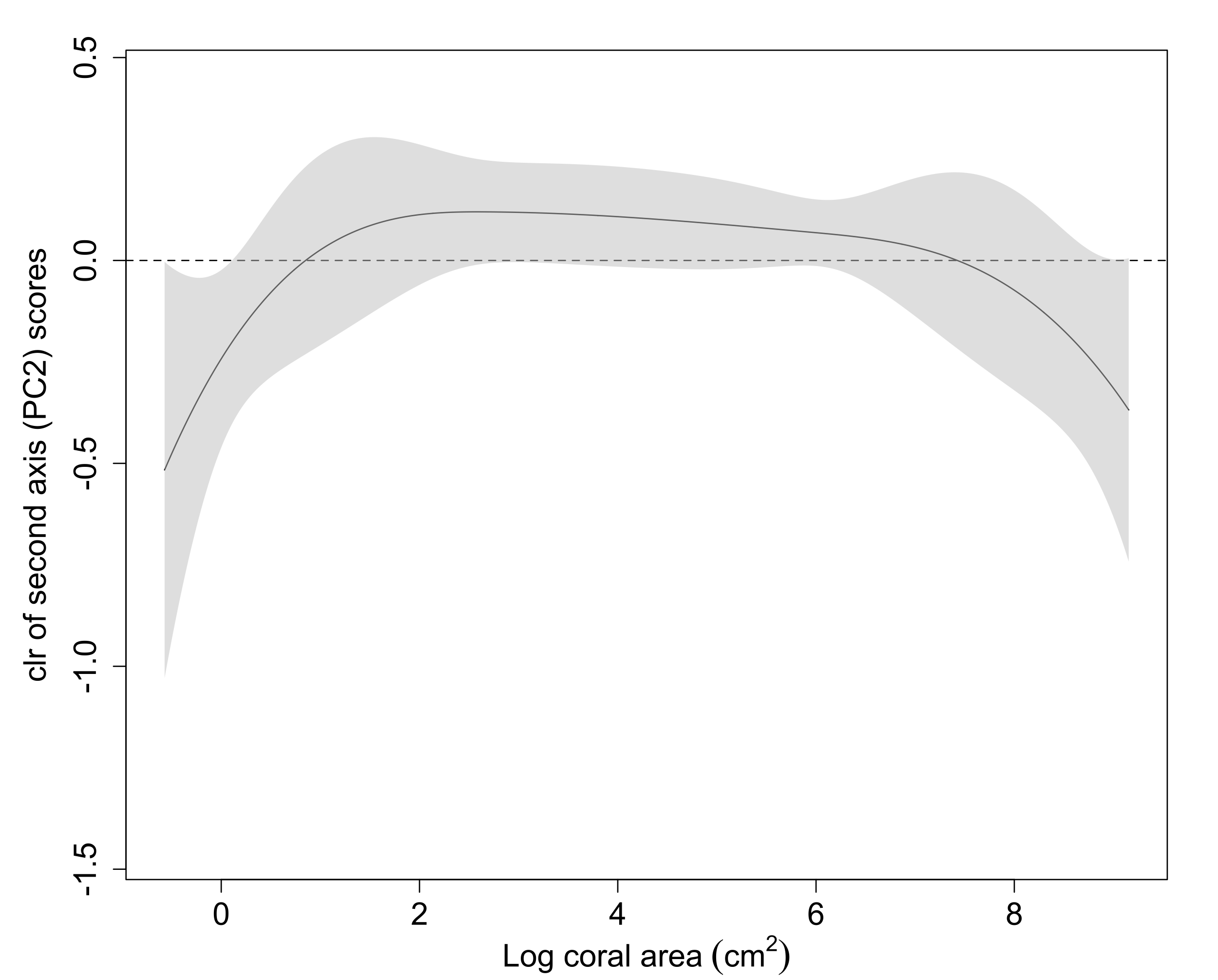

Figure S6. Increases in second axis (PC2) scores might mean lower densities of corals at the extreme sizes but a higher density of corals ~ 1-7 log cm^2^, however no effect is likely as the grey shaded region (asymptotic 95% confidence band) always crossed the dashed line (no effect on the probability densities). Increases in PC2 represent higher minimum chla and kd490 and lower standard deviation of PAR. The black line is the estimated centred log-ratio (clr) transformation of the coefficient function $\boldsymbol{\beta}_{2}$, which measures the effect of a unit increase in PC2 on the probability density of a given log coral area. Positive values on the y-axis mean that the corresponding log coral area on the x-axis becomes more likely as PC2 increases, and negative values mean that the corresponding log coral area becomes less likely.

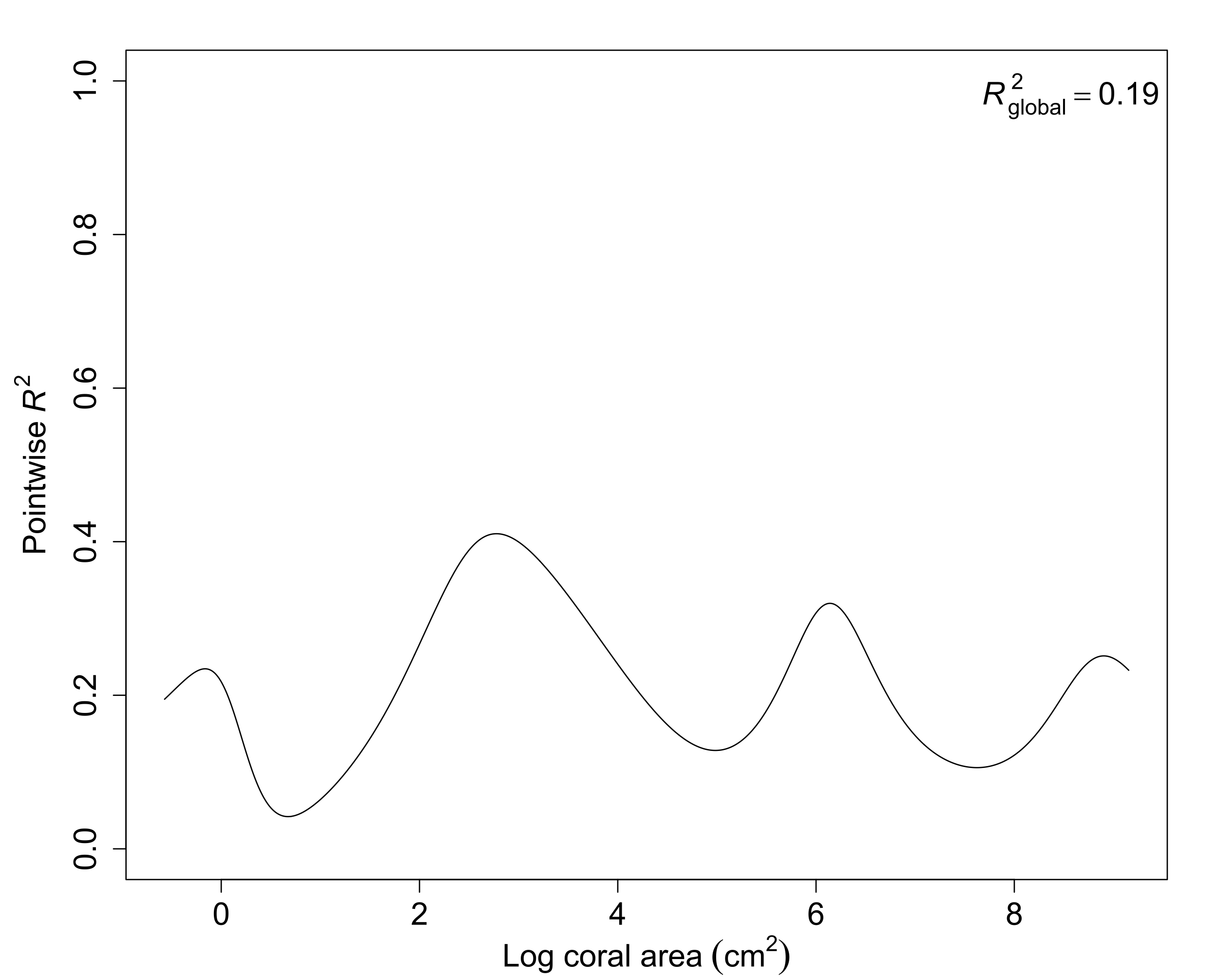
Figure S7. The proportion of variation in probability density explained by the model (pointwise R^2^) at each log coral area, with most explanatory power at approx. 3 log cm^2^, followed by ~6 log cm^2^. Global R^2^ (the total proportion of variation explained by the model) is 0.19.

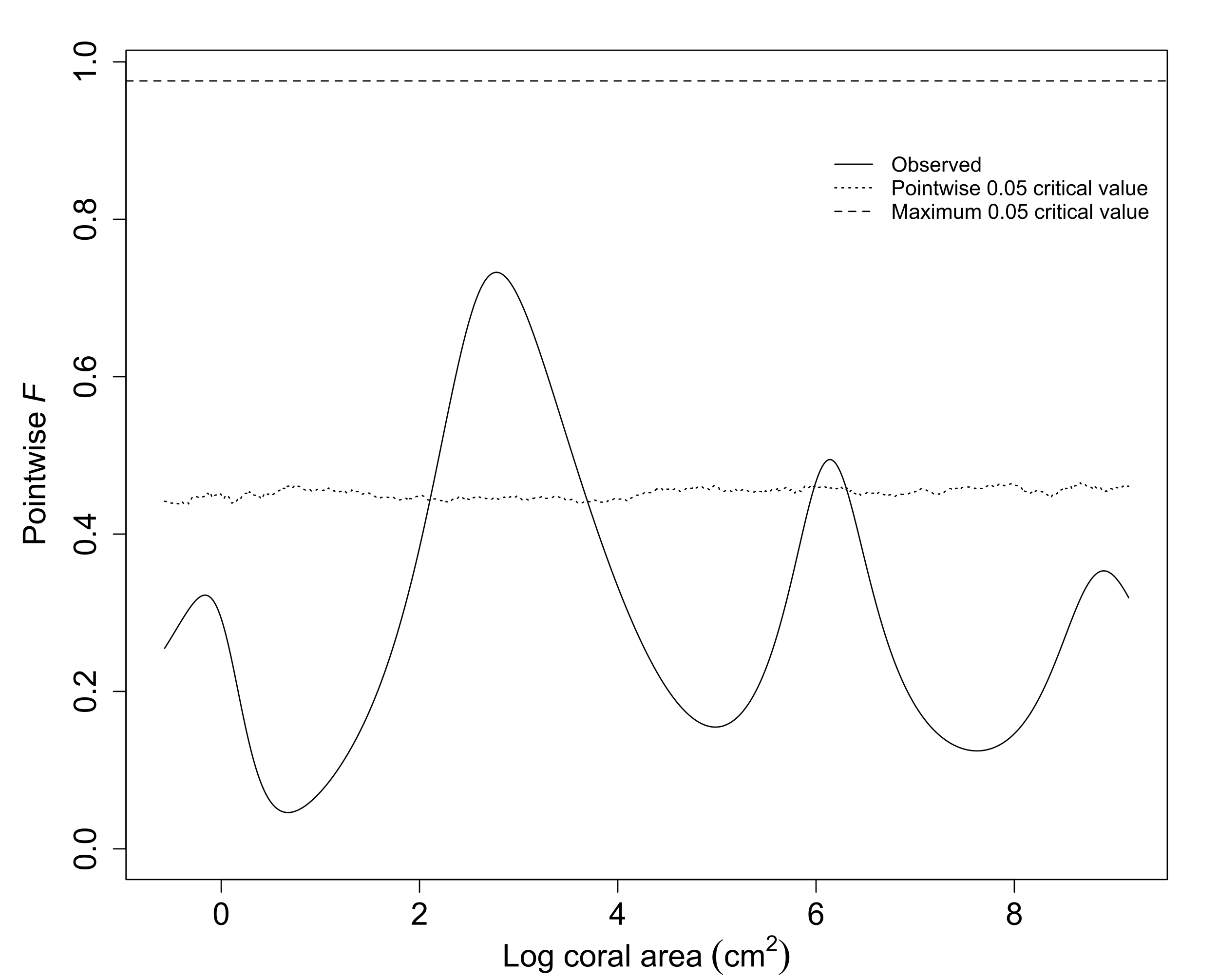

Figure S8. Permutation *F* test for the predictive relationship between log coral area and the PC scores (PC1 and PC2). The observed *F* statistic (solid line) is highest at 3 and 7 log cm^2^, suggesting that the explanatory variables PC1 and PC2 had the most effect there; however, because the observed *F* statistic did not cross the maxima (dashed line) anywhere in the distribution, it was plausible that neither of the explanatory variables affected coral size-frequency distributions. Dotted line: pointwise 0.05 critical values such that only one permuted *F* statistic in 20 exceeds this value at a given log coral area. Dashed line: maximum 0.05 critical value such that only one permuted *F* statistic in 20 exceeds this value at any log coral area. Estimated from 10,000 permutations.

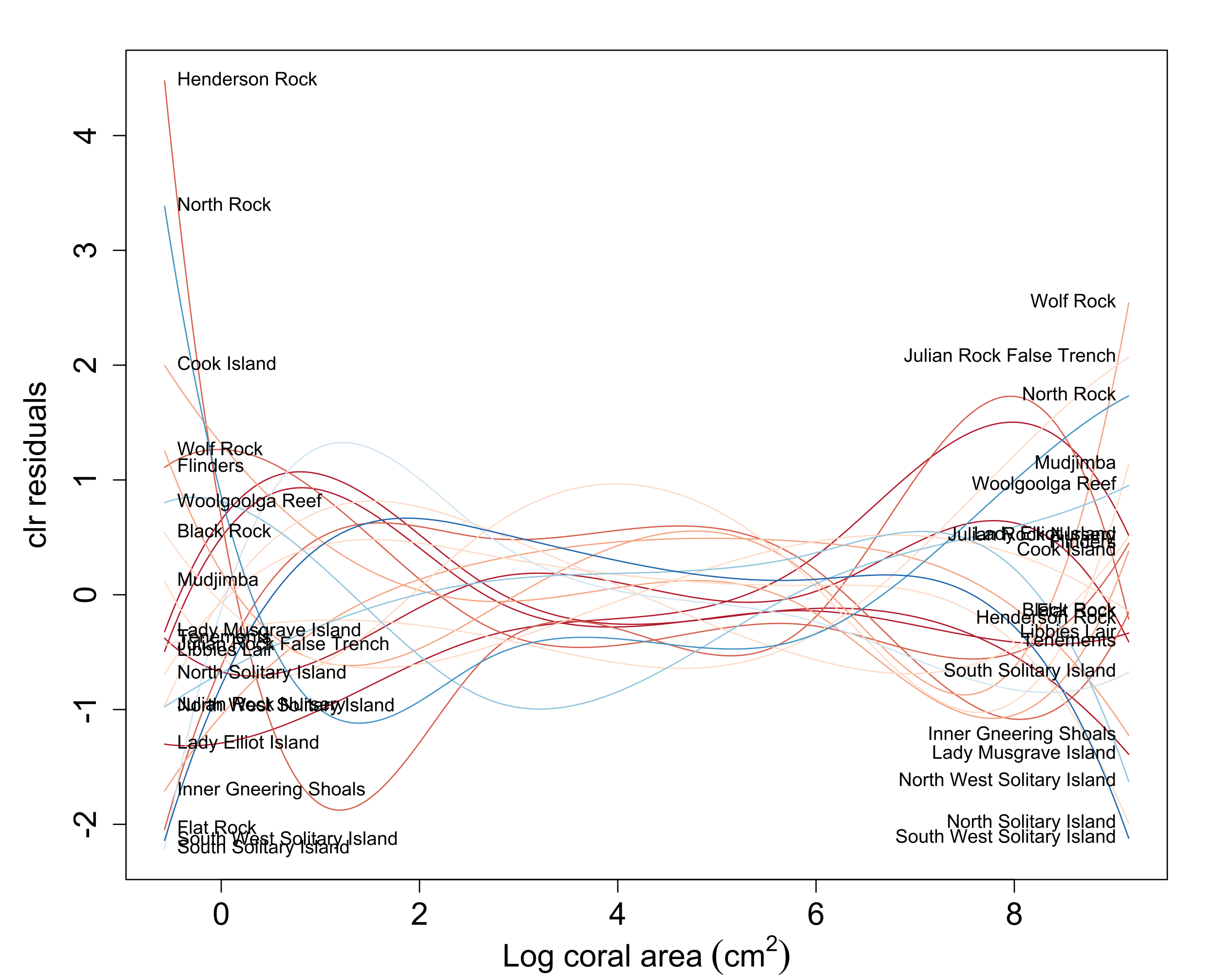

Figure S9. Centred log-ratio (clr) residuals for the predictive relationship between log coral area and the PC scores (PC1 and PC2). Little systematic pattern in the residual functions can be seen, except that they tended to be further from zero at the extreme coral sizes. Line colours go from blue to red in eight steps, where the darkest blue indicates the site with the highest PC1 scores (more turbidity and chlorophyll *a*) and the darkest red are sites with the lowest PC1 scores (high PAR and SST).

### S2: Size-biased sampling

We assume that sampling probability is a function of size alone. This is the case for minus sampling provided that colony shape does not vary systematically with the values of explanatory variables (Baddeley, 1998b, p. 50). Let $g(s)$ be the probability density for sampling colonies of size $s$. Then the size-frequency distribution of sampled colonies, when the true size-frequency distribution has density function $y(s)$, is

$$\frac{y\left( s \right)g(s)}{\int_{a}^{b} y\left( t \right)g\left( t \right)dt}=\boldsymbol{y\oplus g},$$

(Baddeley, 1998a, p. 20). We can show using the change-of-variables formula that the same result holds when the size measure of interest (*e.g.,* log colony area) is a strictly monotone transformation of the size measure that determines sampling bias. Thus, with size-biased sampling, Equation 1 becomes

$$\begin{aligned} \boldsymbol{y}_{\boldsymbol{i}}\boldsymbol{\oplus g=}\boldsymbol{(\beta}_{\boldsymbol{0}}\boldsymbol{\oplus}\left( x_{1,i}\boldsymbol{\odot}\boldsymbol{\beta}_{\boldsymbol{1}} \right)\boldsymbol{\oplus}\left( x_{2,i}\boldsymbol{\odot}\boldsymbol{\beta}_{\boldsymbol{2}} \right)\boldsymbol{\oplus}\boldsymbol{\varepsilon}_{i}\boldsymbol{)\oplus g} \\ \boldsymbol{=}\boldsymbol{(\beta}_{\boldsymbol{0}}\boldsymbol{\oplus g)\oplus}\left( x_{1,i}\boldsymbol{\odot}\boldsymbol{\beta}_{\boldsymbol{1}} \right)\boldsymbol{\oplus}\left( x_{2,i}\boldsymbol{\odot}\boldsymbol{\beta}_{\boldsymbol{2}} \right)\boldsymbol{\oplus}\boldsymbol{\varepsilon}_{i}\boldsymbol{,} \end{aligned}$$

by associativity and commutativity of perturbation. Thus, instead of the true intercept function $\boldsymbol{\beta}_{\boldsymbol{0}}$, we will estimate the perturbed intercept $\boldsymbol{\beta}_{\boldsymbol{0}}\boldsymbol{\oplus g}$, and the overall shape of the estimated response distribution will be biased. However, the estimates of $\boldsymbol{\beta}_{\boldsymbol{1}}$ and $\boldsymbol{\beta}_{\boldsymbol{2}}$, which are of primary interest, will not be affected.
